# Differential control of intestine function by genetically defined enteric neurons

**DOI:** 10.1101/2024.05.02.592229

**Authors:** David Shi, Pranav Reddy, Christopher Walker, McKayla Marrin, Catherine Siu, Nikhil Sharma

## Abstract

The function of the intestine is regulated by direct innervation from a combination of enteric, sensory, and autonomic neurons. A central question in neurobiology is how these distinct peripheral neuron populations collectively control intestinal function. However, disambiguating the functions of intestine-innervating neuronal populations has been a challenge. Using intersectional genetic approaches in mice, we enable precise manipulations of defined neuronal populations within the intestinal tract. We examined enteric neurons, which represent the majority of intestine-innervating neurons, by genetically isolating neuronal subclasses, identifying their morphological specializations, and defining subclass-specific influences on intestinal functions. We further found that food consumption can be modulated by select enteric neuron populations via the spinal sensory afferent pathway. Taken together, the presented molecular genetic characterization of intestine-innervating neurons establishes a foundation for detailed studies of the enteric nervous system and its interactions with the broader neural networks of the body.

## Introduction

The intestine’s ability to perform crucial tasks, such as food transit, waste expulsion, and fluid homeostasis, is governed by its neural circuitry^1-6^. The intestinal tract is innervated by both extrinsic and intrinsic neural populations - the extrinsic composed of the sensory and autonomic neurons, located in distinct ganglia throughout the body, and the intrinsic being the enteric nervous system, which is embedded along the walls of the intestinal tract. Extrinsic neuronal populations connect the intestine to the central nervous system via axonal projections, whereas enteric neurons primarily collateralize within the intestine. Collectively, the enteric, sensory, and autonomic neurons form what could be referred to as the intestinal nervous system. Understanding how these distinct components independently and collaboratively impact intestinal function is a fundamental challenge in neurobiology.

The enteric nervous system comprises the most expansive component of the intestinal nervous system and is estimated to contain an order of magnitude more neurons than all sensory and autonomic ganglia combined^4^. Various methods have been used thus far to categorize the enteric nervous system, namely pharmacological and surgical manipulations, nerve stimulations, immunochemical labeling, and morphological measurements. These efforts have resulted in enteric neuron subpopulations being coarsely catalogued as motor neurons, interneurons, or primary afferent neurons^1-4^, with multiple subclasses existing in each group. In addition, the distinct enteric neuron subclasses have been found to exhibit unique neurochemical marker expression profiles. All together, these form the contemporary neurochemical, functional, and anatomical code that taxonomizes the diversity of neurons in the enteric nervous system^1-7^.

Recent studies have transcriptionally profiled each peripheral neuron lineage (enteric^2,4,8-16^, sensory^10,16-36^, autonomic^7,10,37-42^), defining dozens of transcriptionally distinct populations of neurons in each lineage. Although these studies used different genomic methodologies, it is evident that there is significant cellular heterogeneity in each of these systems and an emergent need to develop approaches to study uncovered subpopulations. While molecular genetic approaches have been crucial in defining functions of neuronal populations across the entire nervous system^25-27,38,43-50^, a major challenge to establishing genetic approaches for the enteric nervous system is the promiscuity of neurochemical markers used to define individual subclasses. For example, many of the marker genes expressed by enteric neurons, such as neuropeptides, can be readily found in extrinsic neuronal populations that also innervate the intestine^2,4,7-42^. This raises the concern that phenotypes, or even morphological features, assigned to specific enteric neurons could potentially be a consequence of inadvertent labeling of other intestine-projecting neuronal populations. We propose that developing sufficiently precise genetic tools will be foundational in enabling rigorous distinction between intrinsic and extrinsic neuronal populations and subsequent investigations into how these systems collaborate to regulate intestine function.

In this study, we developed a molecular genetic approach that allows precise structural and functional examination of the enteric nervous system. We present a compendium of molecular genetic strategies to label a range of enteric neuron subclasses, each innervating distinct structures in the intestine, as well as orthogonal approaches capable of genetically isolating distinct extrinsic neuron lineages. We combine these to implement an intersectional genetic^51^ strategy capable of distinguishing intrinsic and extrinsic neurons. We used these approaches to manipulate neuronal activity in genetically defined enteric neuron subclasses, revealing differential effects on intestinal transit, fecal hydration, and food consumption. Interestingly, we identified subclasses of enteric neurons that require communication with the dorsal root ganglia to regulate food consumption. These findings establish a versatile molecular genetic resource for precise study of the intestinal nervous system. Furthermore, these studies support a model in which enteric neurons not only regulate intestinal function but can also influence feeding behaviors through gut-to-brain pathways defined by enteric-to-dorsal root ganglia signaling.

## Results

### Targeted genetic access to distinguish between peripheral neural lineages

We began by generating a series of mouse lines that could efficiently label all peripheral neurons. We reasoned that this would enable transcriptomic profiling of intestine targeting peripheral neuron lineages with the same sequencing methodology. These markers could then be used to develop mouse models capable of isolating peripheral enteric, sensory, and autonomic neuronal populations. Towards this end, we inserted a recombinase (*FlpO*, *FlpO^ERT2^*, or *Cre^ERT2^*) downstream of the *Elavl4* gene, which has a pan-neuronal expression pattern^52-54^ (**Figure S1**, see methods *’Mouse Generation’* for details). The 2A autocatalytic peptide^55,56^ was inserted between the *Elavl4* gene and the recombinase to minimize disruptions to endogenous gene expression. We then systemically introduced an adeno-associated virus (AAV) carrying a *H2b^mTagGFP2^* expression cassette (CAG:Flp^ON^-H2b^mTagGFP2^) into *Elavl4^T2a-FlpO^* animals to label cells expressing *FlpO*, which restricts labeling to nuclei, rather than axons, thereby facilitating precise localization of marker expression. We harvested the intestine, from the duodenum to the distal colon, from *Elavl4^T2a-FlpO^;CAG:Flp^ON^-H2b^mTagGFP2^* animals and stained tissue sections with antibodies targeting *Phox2b*, a marker of neuronal nuclei in the intestine^57^, and observed robust colocalization with *H2b^mTagGFP2^* in both the myenteric and submucosal plexus (**Figure 1A**). We also observed extrinsic peripheral sensory and autonomic neuronal nuclei were robustly labeled with *H2b^mTagGFP2^*(**Figure 1B**). By contrast, only trace *H2b^mTagGFP2^* signals were detected in non-neuronal cells (**Figure 1F, Row 1**), consistent with the neuronal expression pattern of *Elavl4*. Intestinal tissue from *Elavl4^T2a-CreERT2^;CAG:Cre^ON^-H2b^mTagGFP2^*and *Elavl4^T2a-FlpOERT2^;CAG:Flp^ON^-H2b^mTagGFP2^*also showed robust *H2b^mTagGFP2^* expression restricted to enteric neurons upon tamoxifen dosage and no *H2b^mTagGFP2^* expression in vehicle-treated control animals (**Figure S2**). These measurements confirm that the *Elavl4^T2a-FlpO^* allele broadly labels peripheral neurons, and likewise the *Elavl4^T2a-CreERT2^* and *Elavl4^T2a-FlpOERT2^* alleles label peripheral neurons in a tamoxifen dose-dependent manner.

**Figure 1:**
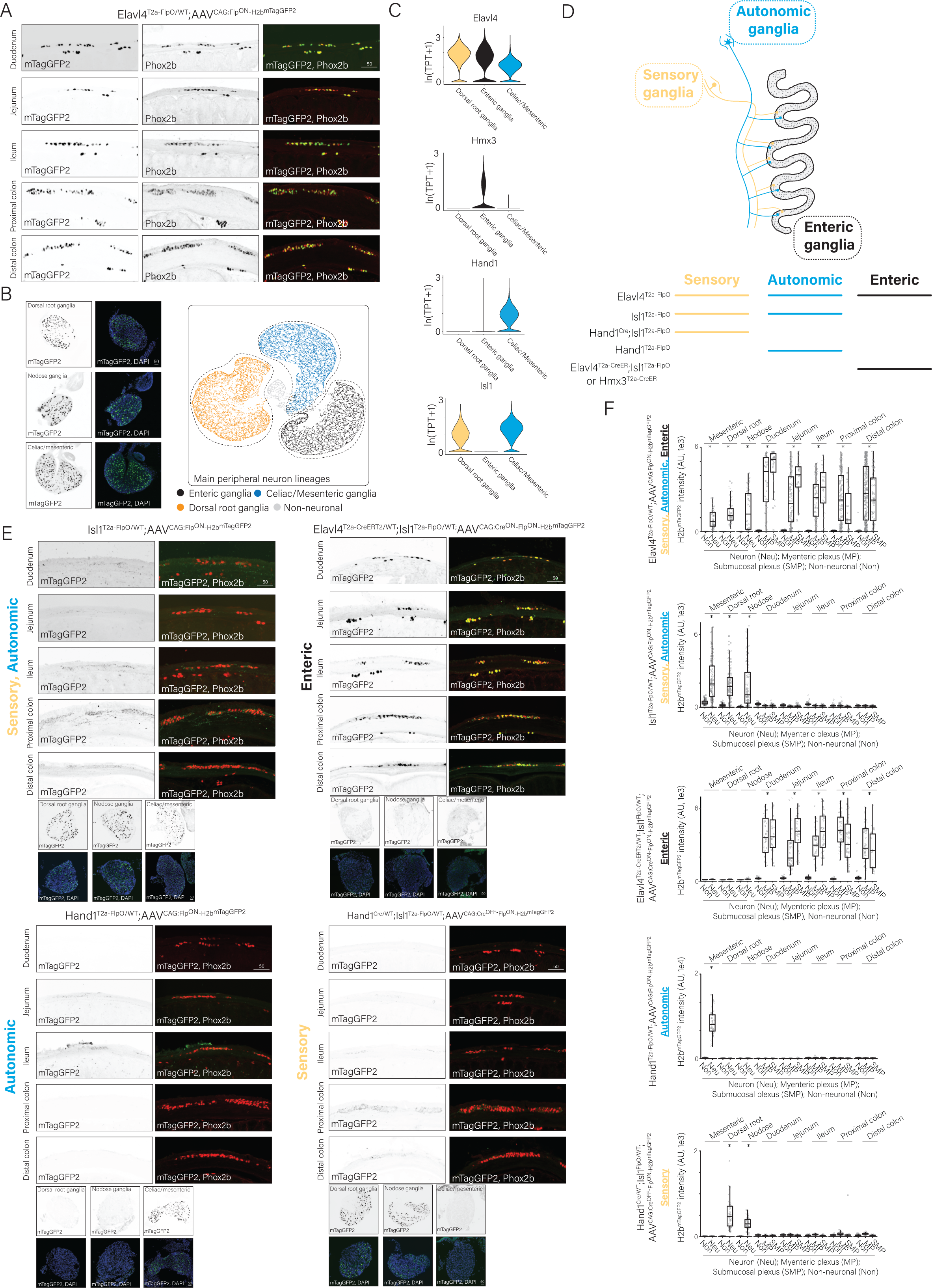
Genetic isolation of peripheral neuron populations that innervate the intestine. **A)** Representative immunostaining images of tissue sections showing colocalization of reporter expression and the neuronal marker gene *Phox2b* in different regions of the intestine from *Elavl4^T2a-FlpO^;CAG:Flp^ON^-H2b^mTagGFP2^*animals. Scale bar = 50µm. **B)** (*Left*) Representative immunostaining images of tissue sections showing *H2b^mTagGFP2^*labeled neurons in the extrinsic ganglia from *Elavl4^T2a-FlpO^;CAG:Flp^ON^-H2b^mTagGFP2^*animals overlaid with and without DAPI. Scale bar = 50µm. (*Right*) UMAP visualizations of scRNA-seq data from enteric, celiac/mesenteric, and dorsal root ganglia. **C)** Violin plots displaying *Elavl4, Hmx3, Hand1,* and *Isl1* expression in the enteric, celiac/mesenteric, and dorsal root ganglia. **D)** (*Top*) Schematic of intrinsic and extrinsic innervation within intestinal tract. (*Bottom*) Schematic illustrating breeding strategies with transgenic mouse lines to isolate particular components of intrinsic or extrinsic innervation. **E)** Representative immunostaining images of tissue sections showing colocalization of reporter expression and the neuronal marker gene *Phox2b* in different regions of the intestine and colocalization of reporter expression and DAPI in the indicated extrinsic ganglia from the same animals from (*Top Left*) *Isl1^T2a-FlpO^;CAG:Cre^ON^-H2b^mTagRFP^*, which labels all extrinsic ganglia (*Top Right*) *Elavl4^T2a-CreERT2^;Isl1^T2a-FlpO^;CAG:Cre^ON^-Flp^ON^-H2b^mTagGFP2^*, which labels all enteric ganglia (*Bottom Left*) *Hand1^T2a-FlpO^;CAG:Flp^ON^-H2b^mTagGFP2^*which labels all peripheral autonomic ganglia (*Bottom Right*) *Hand1^Cre^;Isl1^T2a-FlpO^;CAG:Cre^OFF^-Flp^ON^-H2b^mTagGFP2^*, which labels all sensory ganglia. Scale bar = 50µm. **F)** Quantification of *H2b^mTagXFP^*fluorescence intensity in neuronal and non-neuronal nuclei in extrinsic ganglia (celiac/mesenteric, dorsal root, & nodose) or enteric ganglia in (*Row 1*) *Elavl4^T2a-FlpO^* (*Row 2*) *Isl1^T2a-FlpO^*(*Row 3*) *Elavl4^T2a-CreERT2^;Isl1^T2a-FlpO^* (*Row 4*) *Hand1^T2a-FlpO^* (*Row 5*) *Hand1^Cre^;Isl1^T2a-FlpO^*animals. *p<10^-3^ Wilcoxon-rank sum test compared neuronal and non-neuronal reporter expression. TPT: tags per ten thousand.

We next sought to isolate peripheral ganglia from *Elavl4^FlpO^;CAG:Flp^ON^-H2b^mTagGFP2^*animals for transcriptomic profiling. We separated intact enteric neurons from the large numbers of non-neuronal cells in the intestine by performing fluorescence associated cell sorting (FACS) on resected intestinal tissue (see methods *’Single-cell RNA-sequencing preparation’* for details), and then performed scRNA-seq on isolated enteric neurons to recover single-cell transcriptomes. From this genotype, we extracted the celiac/mesenteric ganglia, which represents a primary source of sympathetic outflow to the intestine, and the dorsal root sensory ganglia. These ganglia were independently subjected to scRNA-seq. Collectively, these efforts generated scRNA-seq datasets under the same platform for the enteric, sensory, and autonomic lineages.

We next compared gene expression profiles between peripheral neuron lineages to identify lineage-restricted marker genes that could serve as drivers for mouse recombinase alleles. To accomplish this, we combined the scRNA-seq datasets from each lineage *in silico* and performed gene expression-based clustering with PCA and visualized these results in two dimensions with UMAP analysis (**Figure 1B, right**). Analysis of the scRNA-seq data revealed that the enteric, autonomic, and sensory ganglia separated into their own distinct individual clusters, and differential gene expression analysis revealed candidate genes with lineage-specific expression profiles (**Figure 1C**). Using these lineage specific candidate markers, we devised strategies to genetically isolate peripheral neurons from the enteric, sensory, or autonomic lineage (**Figure 1D**, see methods *’Mouse Generation’* for details).

We then generated recombinase knock-ins alleles for these candidate genes using a targeting strategy analogous to the *Elavl4* knock-in strategy (**Figure S1**). In select cases, a combination of two marker genes allowed for faithful isolation of the desired neural population (**Figure 1D**). To use ‘intersectional genetic’ strategies, we generated a series of adaptable dual recombinase dependent AAV reporters by taking advantage of a recently generated small self-cleaving type III hammerhead ribozyme variant (T3H48) with traditional recombinase sites^58,59^. In brief, when the T3H48 sequence is in the 3’ UTR of a transgene, it is rapidly destabilized and degraded, thereby preventing protein translation. Flanking the T3H48 ribozyme with recombinase sites stabilizes the transcript in the presence of recombinase. Using this approach, we first confirmed the dual recombinase dependence in heterologous cells (**Figure S3**). We confirmed genetic isolation of extrinsic ganglia by introducing CAG:Flp^ON^-H2b^mTagGFP2^ into *Isl1^T2a-FlpO^* animals (**Figure 1E, Top Left & Figure 1F, Row 2**). We isolated enteric ganglia by introducing CAG:Cre^ON^-H2b^mTagRFP^ into *Hmx3^T2a-CreER^* animals or ‘subtracted’ *Isl1^+^*neurons from *Elavl4^+^* by introducing CAG:Cre^ON^-Flp^OFF^-H2b^mTagGFP2^ into *Elavl4^T2a-CreERT2^;Isl1^T2a-FlpO^* animals. In both genotypes, labeling was restricted to enteric neurons, albeit at modest efficiency in *Hmx3^T2a-CreER^* animals (**Figure 1E, Top Right & Figure 1F, Row 3 & Figure S2C**). We isolate peripheral autonomic ganglia by introducing CAG:Flp^ON^-H2b^mTagGFP2^ into *Hand1^T2a-FlpO^*animals (**Figure 1E, Bottom Left & Figure 1F, Row 4**). Lastly, we isolated sensory ganglia by ‘subtracting’ the *Hand1^+^* autonomic neurons from *Isl1^+^* extrinsic neurons by introducing CAG:Cre^OFF^-Flp^ON^-H2b^mTagGFP2^ into *Hand1^Cre^;Isl1^T2a-FlpO^* animals (**Figure 1E, Bottom Right & Figure 1F, Row 5**). Notably, in each condition we observed little to no labeling of non-neuronal cells (**Figure 1F**). Taken together, these efforts establish a combination of mouse genetic and viral approaches to restrict manipulations to defined neuronal components in the peripheral nervous system.

### Enteric neurons provide dense innervation to all domains of the intestine, whereas extrinsic neurons innervate enteric ganglia

Having established genetic access to each peripheral neuron lineage, we next assayed their independent contributions to innervation of the intestine. To do this, we labeled axons from either all peripheral neurons^Elavl4-T2a-FlpO;CAG:FlpON-GFP^, enteric neurons^Elavl4-T2a-CreERT2;Isl1-T2a-FlpO;CAG:CreON-FlpOFF-GFP^, sensory neurons^Hand1Cre;Isl1-T2a-FlpO;CAG:CreOFF-FlpON-GFP^, or autonomic neurons^Hand1-T2a-FlpO;CAG:FlpON-GFP^ with a cytosolic *GFP*. As expected, labeling all peripheral neurons revealed dense innervation throughout all regions of the intestine, namely the plexuses, circular muscle, and villi (**Figure 2A, right**). Interestingly, we found that the enteric lineage densely innervated the whole intestine (**Figure 2C-D**), whereas the sensory and autonomic lineages most prominently innervated the myenteric and submucosal enteric ganglia, with a comparatively smaller contribution to the circular muscle and villi (**Figure 2E-H**). Taken together, these anatomical measurements indicate that the enteric nervous system provides the largest component of neural innervation to the intestine, with the sensory and autonomic lineages exhibiting densest collateralizations in the enteric ganglia.

**Figure 2:**
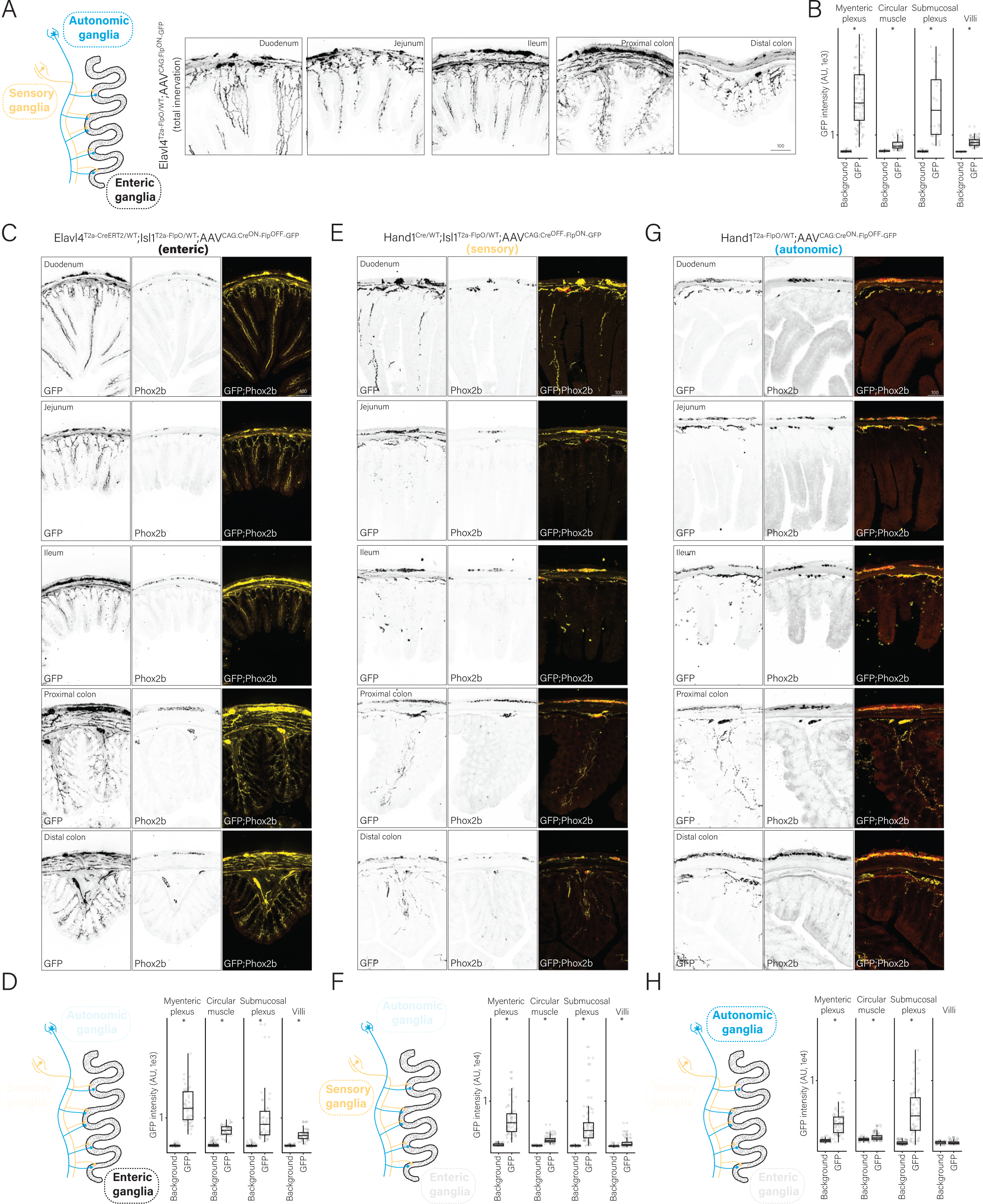
Enteric, sensory, and sympathetic innervation of the intestine. **A)** (*Left*) Schematic of intrinsic and extrinsic innervation of the intestine. (*Right)* Representative immunostaining images of tissue sections showing total innervation in all regions of the intestine from *Elavl4^T2a-FlpO^;CAG:Cre^ON^-*GFP animals. Scale bar = 50µm. **B)** Quantification of fluorescence intensity of axonal GFP in all regions of the intestine from *Elavl4^T2a-FlpO^;CAG:Flp^ON^-GFP*. **C)** Representative immunostaining images of tissue sections showing intrinsic enteric innervation in all regions of the intestine from *Elavl4^T2a-CreERT2^;Isl1^T2a-FlpO^-CAG:Cre^ON^-Flp^OFF^-GFP* animals. Scale bar = 100µm. **D)** Quantification of fluorescence intensity of axonal GFP in all regions of the intestine from *Elavl4^T2a-CreERT2^;Isl1^FlpO^-CAG:Cre^ON^-Flp^OFF^-GFP* animals. **E)** Representative immunostaining images of tissue sections showing extrinsic sensory innervation in all regions of the intestine from *Hand1^T2a-Cre^;Isl1^T2a-FlpO^;CAG:Cre^OFF^-Flp^ON^-GFP* animals. Scale bar = 100µm. **F)** Quantification of fluorescence intensity of axonal GFP in all regions of the intestine from *Hand1^T2a-Cre^;Isl1^T2a-FlpO^;CAG:Cre^OFF^-Flp^ON^-GFP* animals. **G)** Representative immunostaining images of tissue sections showing extrinsic autonomic innervation in all regions of the intestine from *Hand1^T2a-FlpO^;CAG:Cre^ON^-Flp^OFF^-GFP* animals. Scale bar = 100µm. **H)** Quantification of fluorescence intensity of axonal GFP in all regions of the intestine from *Hand1^T2a-FlpO^;CAG:Cre^ON^-Flp^OFF^-GFP* animals. *p<10^-3^ Wilcoxon-rank sum test.

### Curation and generation of genetic approaches for enteric neuron subclasses

Given that the enteric nervous system presented as the most prominent contributor to intestinal innervation, we next began establishing genetic strategies to access distinct subclasses of enteric neurons for subsequent morphological and functional analyses. We first clustered and visualized our scRNA-seq transcriptomes recovered from enteric neurons by PCA/UMAP analysis (**Figure S4A-C**). Each cluster was taxonomized according to the frequency of cells recovered by scRNA-seq, with the largest cluster by cell count being denoted by the Greek letter α. Consistent with previous scRNA-seq taxonomic studies of the enteric nervous system^8,10,11^, our analysis revealed a similar range of transcriptionally distinct subclasses of enteric neurons (**Figure S4**). To further confirm that single cells recovered from the *Elavl4^T2a-FlpO^*allele aligned with previously published scRNA-seq datasets, we performed FACS isolation of intact enteric neurons using the previously used^11^ *Actl6b^Cre^* (also known as Baf53b-Cre^60^) allele crossed to a Cre-dependent *H2b^mTagRFP^*fusion that we knocked into the Rosa26 locus (*Rosa26^LSL-H2b-mTagRFP^*) (**Figure S1E**). We observed that single cells recovered from these animals revealed similar clustering profiles to enteric neurons recovered from the *Elavl4^T2a-FlpO^* allele (**Figure S4E-I**).

We next identified subclass-specific marker genes and used them to create knock-in lines that would provide genetic access to these populations (**Figure S4A-C**). Towards this end, we obtained the *Oprk1^T2a-Cre^*allele^17^ and generated the *Kcns3^T2a-Cre^* allele for the α subclass; for the β subclass we generated the *Cox8b^T2a-FlpO^*allele; for the γ subclass we obtained the *Cck^IRES-Cre^* allele^43^; for the δ subclass we obtained the *Gad2^IRES-Cre^* allele^43^; for the ε subclass we generated the *Gucy2g^T2a-Cre^* allele; for the ζ subclass we generated the *Nxph2^T2a-Cre^* allele; for the η subclass we generated the *Cysltr2^T2a-FlpO^* allele and obtained the *Cysltr2^T2a-Cre^* allele^17^; and for the θ subclass we obtained the *Sst^IRES-Cre^* allele^43^ (**Figure S1**). We were unable to obtain successful germline transmission events for the *Gucy2g^T2a-Cre^* allele despite multiple attempts, and therefore were unable to include this subclass in subsequent analyses. We initially characterized the specificity of each driver line in the enteric nervous system by introducing a *GFP* reporter via AAV transduction into each mouse line and performing double smRNA-FISH analyses for *GFP* and the target gene used to drive recombinase expression. These analyses confirmed that a highly faithful and efficient genetic strategy was identified for the α, β, γ, δ, ζ, θ, and η enteric neuron subclasses (**Figure 3**, see methods *’Mouse Generation’* for details). Taken together, these efforts present a series of mouse alleles that label transcriptionally distinct subclasses of enteric neurons.

**Figure 3.**
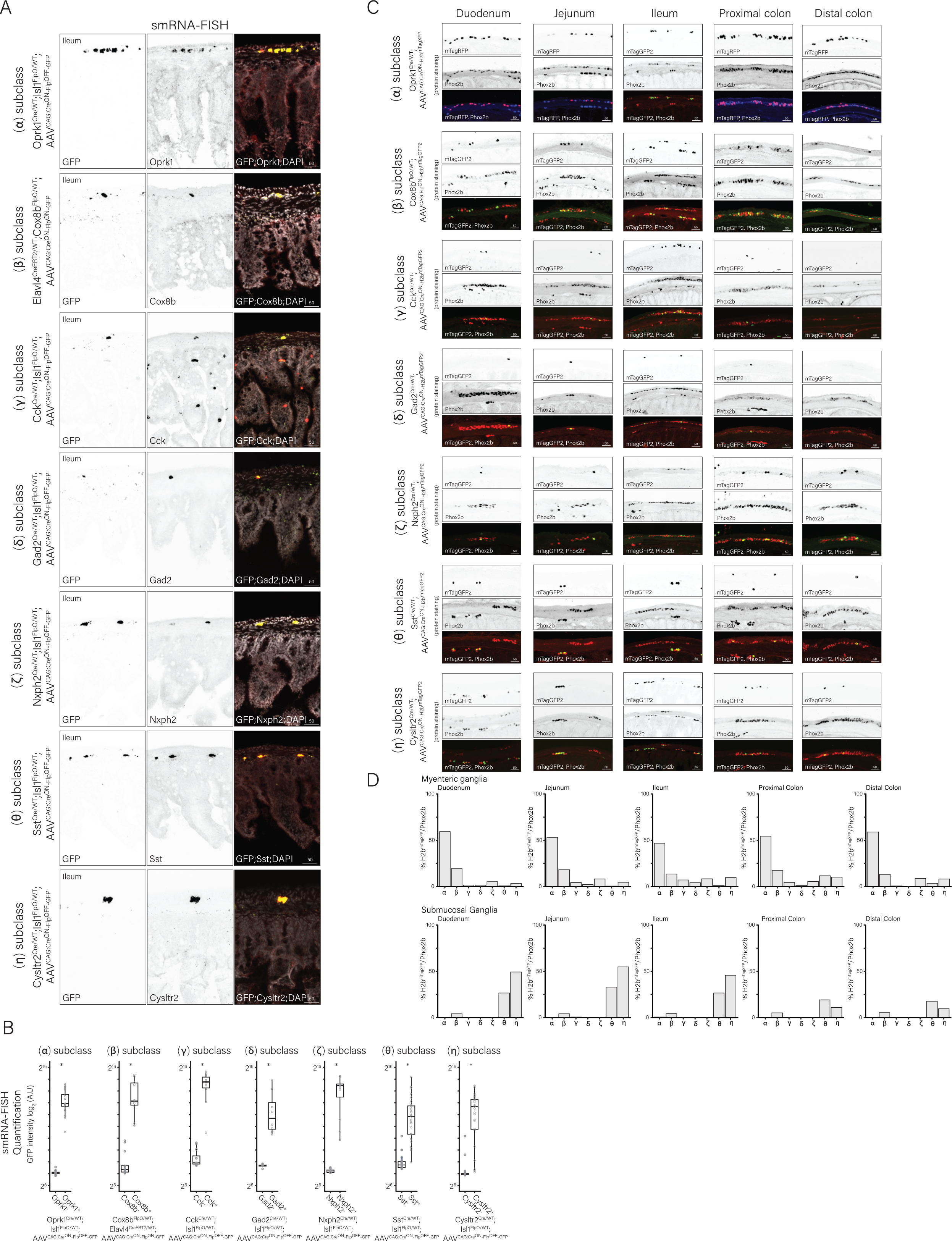
Development of genetic tools for, and localization, of enteric neuron subclasses. **A)** Genetic strategies to label enteric neuron subclasses were transduced with an AAV carrying a recombinase dependent *GFP*. Representative images of double smRNA-FISH analysis in ileum sections with probes targeting *GFP* and the endogenous gene driving the recombinase for the (*Row 1*) α subclass ^(*Oprk1-T2a-Cre;Isl1-T2a-FlpO*)^; (*Row 2*) β subclass ^(*Elavl4-T2a-CreER;Cox8b-T2a-FlpO)*^; (*Row 3*) γ subclass ^(*Cck-IRES-Cre;Isl1-T2a-FlpO*)^; (*Row 4*) δ subclass ^(*Gad2-IRES-Cre;Isl1-T2a-FlpO*)^; (*Row 5*) ζ subclass ^(*Nxph2-T2a-Cre;Isl1-T2a-FlpO*)^; (*Row 6*) θ subclass ^(*Sst-IRES-Cre;Isl1-T2a-FlpO*)^; (*Row 7*) η subclass ^(*Cysltr2-T2a-Cre;Isl1-T2a-FlpO*)^. **B)** Quantification GFP signal intensity in targeted neurons from each subclass compared to neighboring neurons not expressing the recombinase. **C)** Representative confocal microscopy images of *H2b^mTagXFP^* to reveal nuclei, the neuronal marker *Phox2b*, and an overlay in the (*Left to Right*) duodenum to distal colon from (*Row 1*) α subclass ^(*Oprk1-T2a-Cre;Isl1-T2a-FlpO*)^; (*Row 2*) β subclass ^(*Elavl4-T2a-CreER;Cox8b-T2a-FlpO)*^; (*Row 3*) γ subclass ^(*Cck-IRES-Cre;Isl1-T2a-FlpO*)^; (*Row 4*) δ subclass ^(*Gad2-IRES-Cre;Isl1-T2a-FlpO*)^; (*Row 5*) ζ subclass ^(*Nxph2-T2a-Cre;Isl1-T2a-FlpO*)^; (*Row 6*) θ subclass ^(*Sst-IRES-Cre;Isl1-T2a-FlpO*)^; (*Row 7*) η subclass ^(*Cysltr2-T2a-Cre;Isl1-T2a-FlpO*)^. **D)** Bar plots quantifying the fraction of enteric neurons labeled by each recombinase driver line in each segment of intestine in the myenteric or submucosal plexus. *p<10^-3^ Wilcoxon-rank sum test.

### Genetically defined enteric neurons differentially modulate intestinal function in vivo

We next aimed to assess the contribution of the genetically defined enteric neuron subclasses on intestine function in vivo. However, we noted that the markers used to develop mouse lines that distinguish neurons of the enteric nervous system were often found to also be expressed in extrinsic neuronal populations (**Figure S4D**). Therefore, to ensure that our genetic manipulations were restricted to the enteric nervous system, we injected a Cre^ON^-Flp^OFF^ AAV encoding the excitatory designer receptor *hM3Dq*^61^ into mice with the general genotype *Subclass^Cre^;Isl1^T2a-FlpO^*. In these animals, Cre expression would allow for enteric subclass-specific labeling, while extrinsic neuronal populations express FlpO and therefore would have no *hM3Dq* expression. Upon intraperitoneal injection of clozapine-N-oxide (CNO), which is inert in wild-type animals (**Figure S5**), neurons expressing the *hM3Dq* receptor would transiently increase their firing rates^61^. Using these animals, we measured intestinal transit non-invasively by tracking fecal production and examined the hydration content of fecal pellets (**Figure S5,** see methods *’Behavioral measurements’* for details). We found that CNO-dependent activation increased detected fecal event rates in the α, β, ζ, θ, and η enteric neuron subclasses compared to control groups, which included mice of the same genotypes injected with vehicle only (**Figure 4A**). By contrast, we found that CNO-dependent activation of the γ and δ enteric neuron sub-classes led to a decrease in detected fecal events compared to vehicle injected controls (**Figure 4A**). Notably, fecal hydration was also affected in a subset of enteric neuron subclasses upon activation with CNO, with the β, θ, and η populations leading to an increase in fecal hydration content, whereas the γ and δ subclasses showed a subtle decrease in fecal hydration compared to vehicle-treated control animals (**Figure 4B**). These studies suggest that each of these genetically defined populations have unique effects on intestinal transit and fecal hydration.

**Figure 4:**
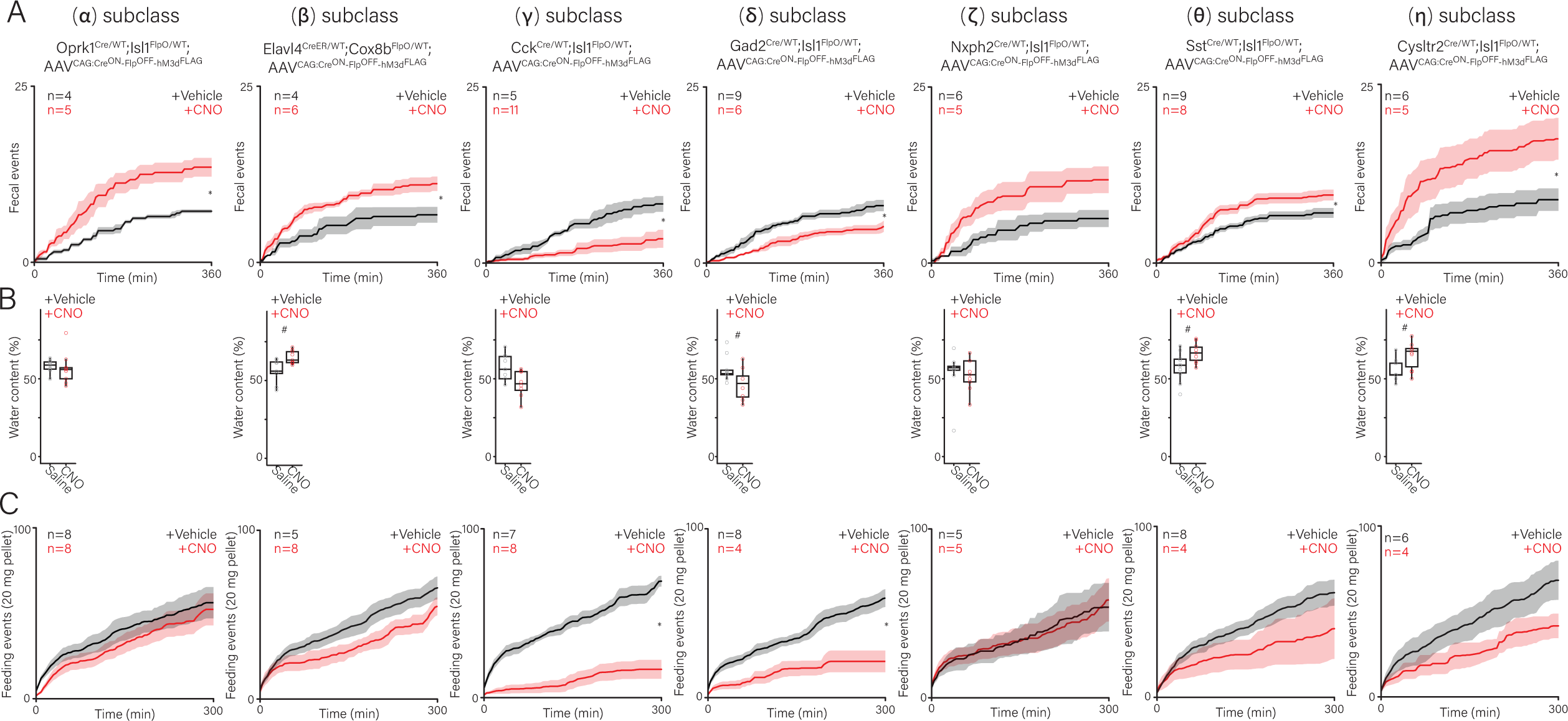
Enteric neuron activation reveals subclass specific effects on intestinal transit, fecal hydration, and food consumption. **A)** Cumulative fecal production over six hours observed in response to saline (vehicle) or CNO injected treatment in the indicated genotypes. Black traces represent the means for vehicle treated animals whereas red traces represent the means for CNO treated animals. Shaded areas represent mean ± SEM. **B)** Stool samples were collected and weighed immediately after extrusion and again after completely dried. Individual data points represent percentage of water weight for single stool samples from the indicated genotype and condition. The bars represent median ± IQR. **C)** Cumulative 20 mg food pellet acquisition events recorded in overnight fasted animals treated with vehicle or CNO over six hours in the indicated genotypes. Black traces represent the means for vehicle treated animals whereas red traces represent the means for CNO treated animals. Shaded areas represent mean ± SEM. In all cases, from left to right, the genotypes display are α subclass ^(*Oprk1-T2a-Cre;Isl1-T2a-FlpO*)^; β subclass ^(*Elavl4-T2a-CreER;Cox8b-T2a-FlpO*)^; γ subclass ^(*Cck-IRES-Cre;Isl1-T2a-FlpO*)^; δ subclass^(*Gad2-IRES-Cre;Isl1-T2a-FlpO*)^; ζ subclass ^(*Nxph2-T2a-Cre;Isl1-T2a-FlpO*)^; θ subclass ^(*Sst-IRES-Cre;Isl1-T2a-FlpO*)^; η subclass ^(*Cysltr2-T2a-Cre;Isl1-T2a-FlpO*)^. ^#^p<10^-2^ two way t-test. *p<0.05 Wilcoxon-rank sum test done on total cumulative fecal pellets and cumulative food pellets retrieved, comparing vehicle or CNO injected animals.

We further analyzed whether activation of the genetically defined enteric neuron subclasses would affect food consumption. We fasted mice overnight and then exposed them to food, quantifying the amount of food acquired by the animals upon reintroduction of food (**Figure S5,** see methods *’Behavioral measurements’* for details). By injecting mice with CNO or vehicle control during the re-exposure to food, we could assess if the activation of an enteric neuron subclass would alter food consumption. Interestingly, we observed that after fasting and upon CNO treatment, the γ and δ enteric neuron subclasses led to a robust reduction in food consumption compared to saline controls, whereas the other subclasses exhibited modest to no reduction in food consumption (**Figure 4C**). We noted that activation of enteric neuron subclasses that led to a decrease in fecal production also showed the greatest reduction in post-fasting food acquisition. Taken together, these data indicate that enteric neuron subclasses can exert distinct forms of modulation on intestinal transit, fecal hydration, and food consumption.

### Genetic labeling of enteric neuron subclasses reveals distinct innervation patterns

Having identified how the genetically defined enteric neuron subclasses influence intestinal function, we next aimed to define features that could facilitate these functions. To begin, we hypothesized that the axonal innervation patterns of enteric neurons, as with all other neurons, would be related to their functionality. We further reasoned that neuronal morphology, when combined with gene expression profiles and effects on intestinal function, could aid in developing mechanistic models for how each enteric neuron subclass modulates intestine function. Towards this end, we transduced animals (e.g., *Subclass^Cre^;Isl1^T2a-FlpO^*) with a cytosolic *GFP* reporter using an AAV carrying a CAG:Cre^ON^-Flp^OFF^-GFP expression cassette to ensure *GFP* expression was indeed restricted to the Cre-expressing enteric neuron subclass.

Analyzing GFP labeling from these animals, we found that axons of the α^(*Oprk1-T2a-Cre;Isl1-T2a-FlpO*)^ and β^(*Elavl4-*^ *^T2a-CreER;Cox8b-T2a-FlpO^*^)^ enteric neuron subclass primarily innervated the circular muscle (**Figure 5A,B**); axons of the γ^(*Cck-IRES-Cre;Isl1-T2a-FlpO*)^, δ^(*Gad2-IRES-Cre;Isl1-T2a-FlpO*)^, and ζ^(*Nxph2-T2a-Cre;Isl1-T2a-FlpO*)^ subclasses appeared to aggregate around other enteric ganglia (**Figure 5C-E**); axons of the θ^(*Sst-IRES-Cre;Isl1-T2a-FlpO*)^ subclass primarily sent projections to the intestinal crypts and villi (**Figure 5F**); and lastly, axons of the η^(*Cysltr2-T2a-Cre;Isl1-T2a-*^ *^FlpO^*^)^ subclass densely labeled the villi and displayed robust axonal signal in other enteric ganglia with axons that appeared to cross the circular muscle en route to myenteric ganglia (**Figure 5G**). We note the recovered profiles of each genetically defined subclass were largely similar from the duodenum to the distal colon (**Figure S6, S7**), albeit with slight variations. For example, the β enteric neuron subclass exhibited detectable *GFP* signal in the villi particularly in the duodenum; however, this signal was significantly less than what was observed in the circular muscle.

**Figure 5:**
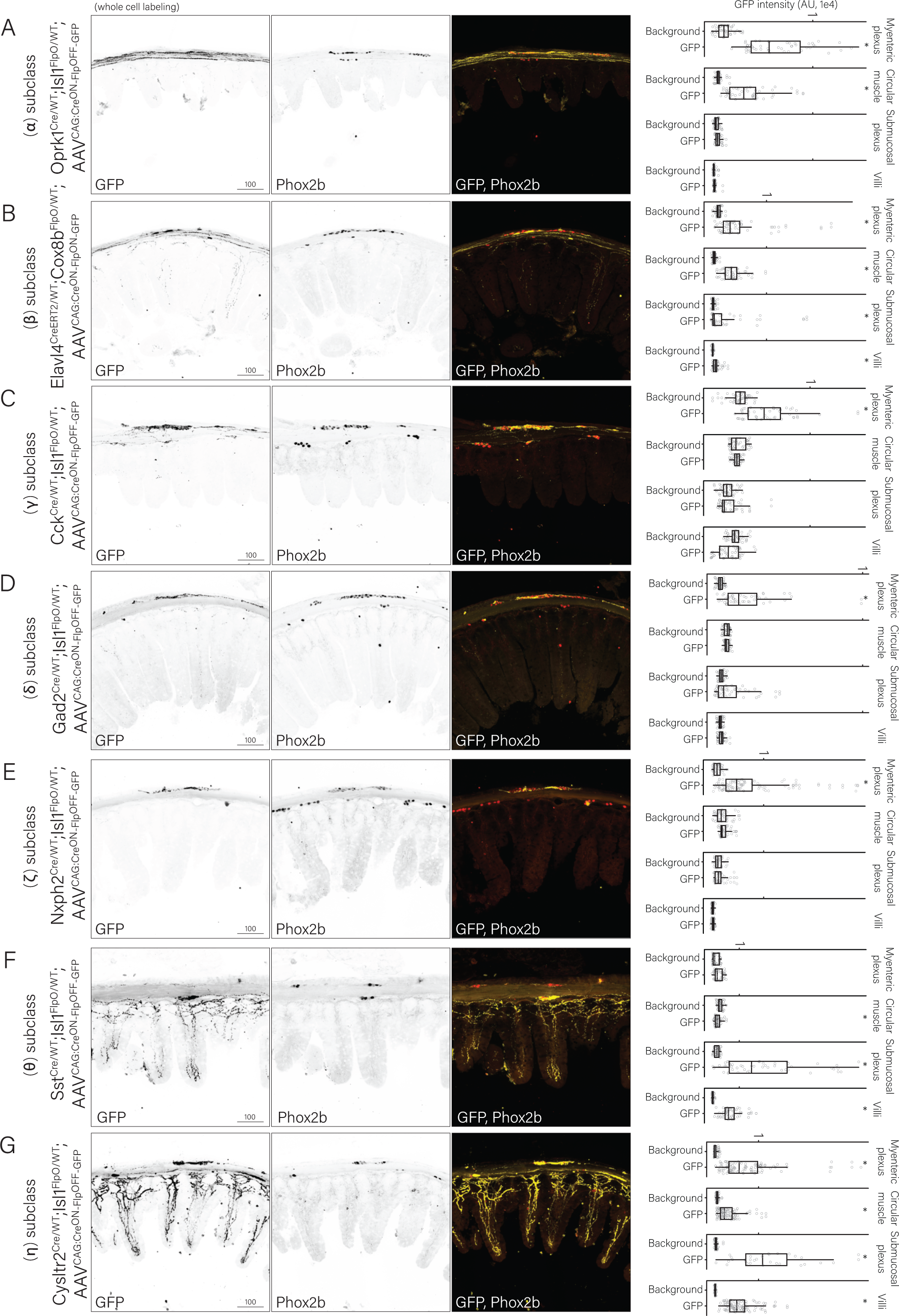
The genetically defined enteric neuron subclasses display distinct morphological profiles in the intestine. **A)** Representative confocal image of ileal tissue sections in which enteric α subclass ^(*Oprk1-T2a-*^ *^Cre;Isl1-T2a-FlpO^*^)^ is labeled with cytosolic GFP to reveal axonal arbors. **B)** Representative confocal image of ileal tissue sections in which enteric β subclass ^(*Elavl4-T2a-*^ *^CreER;Cox8b-T2a-FlpO^* is labeled with cytosolic GFP to reveal axonal arbors. **C)** Representative confocal image of ileal tissue sections in which enteric γ subclass ^(*Cck-IRES-Cre;Isl1-*^ *^T2a-FlpO^*^)^ is labeled with cytosolic GFP to reveal axonal arbors. **D)** Representative confocal image of ileal tissue sections in which enteric δ subclass ^(*Gad2-IRES-*^ *^Cre;Isl1-T2a-FlpO^*^)^ is labeled with cytosolic GFP to reveal axonal arbors. **E)** Representative confocal image of ileal tissue sections in which enteric ζ subclass ^(*Nxph2-T2a-Cre;Isl1-*^ *^T2a-FlpO^*^)^ is labeled with cytosolic GFP to reveal axonal arbors. **F)** Representative confocal image of ileal tissue sections in which enteric θ subclass ^(*Sst-IRES-Cre;Isl1-*^ *^T2a-FlpO^*^)^ is labeled with cytosolic GFP to reveal axonal arbors. **G)** Representative confocal image of ileal tissue sections in which enteric η subclass ^(*Cysltr2-T2a-*^ *^Cre;Isl1-T2a-FlpO^*^)^ is labeled with cytosolic GFP to reveal axonal arbors. **(A-G)** *(Left)* Representative image of cytosolic *GFP*, the neuronal marker *Phox2b*, and an overlay in the ileum, *(Right)* Quantification of fluorescence intensity of cytosolic *GFP* in the indicated location. *p<10^-3^ Wilcoxon-rank sum test. Scale bar for cytosolic GFP/Phox2b images= 100 µm. GFP intensities for all images were in the range of 10^4^ except for enteric γ subclass ^(*Cck-IRES-Cre;Isl1-T2a-*^ *^FlpO^*^)^, which was in the range of 10^3^.

We further determined the proportion of the total enteric neuron population represented by each genetically defined subclass by introducing a recombinase-dependent *H2b^mTagXFP^* reporter, which allowed for quantitation of neuronal nuclei (**Figure 3C,D**). These analyses revealed that the α, β, θ, η subclasses were found across the duodenum to the distal colon (**Figure 3C,D**), with the α and β neurons collectively representing nearly 75% of all enteric neurons. By contrast, we found that the γ and δ subclasses were preferentially enriched in the ileal and proximal colon segments of the intestine (**Figure 3C,D**), and were comparatively rare, representing ∼5% of all enteric neurons. Interestingly, we noted that β subclass was found in both the myenteric and submucosal plexus, while several subclasses were enriched primarily in the myenteric plexus (α, ζ). By contrast, the θ and η subclasses were enriched in the submucosal plexus. We further confirmed this localization preference by performing scRNA-seq on FACS-isolated intact whole cells from submucosal plexuses acutely dissected and dissociated from ileal segments of *Elavl4^T2a-FlpO^;CAG:Flp^ON^-H2b^mTagGFP2^*animals (**Figure S8A**). Taken together, these analyses indicate that the genetically defined enteric neuron subclasses encompass a remarkable range of morphological diversity.

### Inferring morphological and functional features from gene expression patterns in genetically defined enteric neuron subclasses

We next sought to further characterize the genetically defined populations by examining their neurochemical and neurotransmitter profiles. We assigned the α, γ, ζ, θ, and η subclasses as cholinergic neurons based on their expression of markers of the excitatory neurotransmitter acetylcholine (*Chat*, *Slc18a3*); similarly, we categorized the β and δ subclasses as nitrergic based on the expression of genes involved in nitric oxide biosynthesis (*Nos1*) (**Figure S8B**). We confirmed the validity of these assignments by performing scRNA-seq with FACS-isolated intact whole cells from recombinase driver lines under the control of *Chat*^62^ or *Nos1*^63^. Consistent with α, γ, ζ, θ, and η subclasses being of cholinergic identity, these subclasses were recovered when we FACS isolated intact single cells and performed scRNA-seq analysis of enteric neurons from *Chat^IRES-Cre^;CAG:Cre^ON^-H2b^mTagGFP2^*mice (**Figure S8C, top**). The β and δ subclasses, which do not express *Chat* and therefore are unlikely to be of cholinergic identity, were not recovered in *Chat^IRES-Cre^;CAG:Cre^ON^-H2b^mTagGFP2^*FACS isolations. Instead, they were recovered in scRNA-seq analysis from FACS-isolated intact single cells obtained from *Nos1^IRES-CreER^;CAG:Cre^ON^-H2b^mTagGFP2^* animals, consistent with the gene expression data indicating that they were nitrergic (**Figure S8C, bottom**). Based on the gene expression, morphological, and physiological observations, we propose that the α and β subclasses contain circular muscle targeting neurons, with the α subclass being cholinergic and the β subclass being nitrergic; the γ, δ, and ζ subclasses contain enteric interneurons with the γ and ζ being cholinergic and the δ being nitrergic; the θ subclass contains cholinergic neurons with an anatomical pattern consistent with an intrinsic primary afferent identity; and the η subclass contains cholinergic neurons with secretomotor neuron functionality.

From the axonal innervation pattern, gene expression profile, and physiological function, we observed that the largest populations of genetically defined enteric neuron subclasses (α and β) primarily innervated the circular smooth muscle along the intestine. We further assessed that the α population was cholinergic and β population nitrergic (**Figure S8**). The activation of these populations resulting in elevated intestinal transit and fecal production is consistent with the ‘law of the intestine’ proposed by Startling and Bayliss in 1899, stating that directional movement of intestinal content is due to oral excitation and aboral relaxation^64-66^. A hypothesized cellular mechanism based on ex vivo DiI tracing^67-72^ from excised intestine tissue would be that the excitatory α neurons project orally, while inhibitory β neurons project aborally. To test whether the α/β populations indeed have polarized axonal projection patterns, we sought to reconstruct the axonal arbors of individual neurons using a titratable intersectional genetic strategy (α^Oprk1-T2a-Cre;Elavl4-T2a-FlpERT2^ and β^Cox8b-T2a-FlpO;Elavl4-T2a-CreERT2^). Towards this end, we introduced a dual recombinase-dependent alkaline phosphatase (AP)^73^ reporter (CAG:Cre^ON^-Flp^ON^-AP) into a cross where sufficiently low doses of tamoxifen could be used to titrate the number of reporter-positive neurons to a small enough number to allow for reconstruction of single neuron axonal arbors. Strikingly, we observed that neurons from the α subclass had axonal projections that initially extended in the oral direction and then subsequently bifurcated into the circular muscle (**Figure S9A, C**). By contrast, neurons of the β subclass initially extended in the aboral direction prior to similarly bifurcating into the circular muscle (**Figure S9B, C**). These results support a model in which the α and β subclasses label excitatory and inhibitory smooth muscle motor neurons. Taken together, we propose more broadly that a combination of behavioral, anatomical, and gene expression profiles allow for putative functional assignments to be mapped onto the genetically defined enteric neuron subclasses.

### An enteric to dorsal root ganglia signal modulates food intake but not intestinal transit

How might enteric neurons affect food consummatory behaviors? We observed that enteric neuron activation not only led to changes in intestinal transit or fecal hydration, but that select subclasses (γ and δ) also displayed reductions in post-fasting food intake. We further observed that the γ and δ subclasses were enriched in the ileal and proximal colon segments of the intestinal tract. Given the documented roles of the brain in food consumption behaviors^74^, we hypothesized that food intake suppression by enteric neurons would involve communication with the central nervous system via extrinsic sensory pathways that innervate the intestine. However, we note that there are two distinct sensory pathways innervating the intestine, one mediated by nodose ganglia sensory neurons and the other by the dorsal root ganglia sensory neurons.

We first aimed to determine the relative contributions of sensory neuron innervation, either from the nodose or dorsal root ganglia, to the ileum and proximal colon, thereby informing which would serve as a candidate for γ and δ enteric neuron-mediated food intake suppression. To distinguish between innervation from these two sensory pathways, we first crossed the *Hand1^Cre^*allele^75^, which labeled the celiac/mesenteric ganglia but not the dorsal root or nodose ganglia, to the *Isl1^T2a-FlpO^* allele. In *Hand1^Cre^;Isl1^T2a-FlpO^* animals, we ensured viral delivery of reporter would not label any peripheral autonomic ganglia as they could be excluded due to their expression of Cre recombinase. We could then further distinguish sensory neurons of the nodose from dorsal root ganglia by directly injecting AAV vectors into each respective ganglia (**Figure 6A-F**). We found that the ileal and proximal colon segments showed a higher density of innervation from the dorsal root ganglia in comparison to the nodose ganglia (**Figure 6A-F**). These results suggest that the dorsal root ganglia may serve as the sensory pathway that mediates γ and δ enteric neuron food intake suppression.

**Figure 6:**
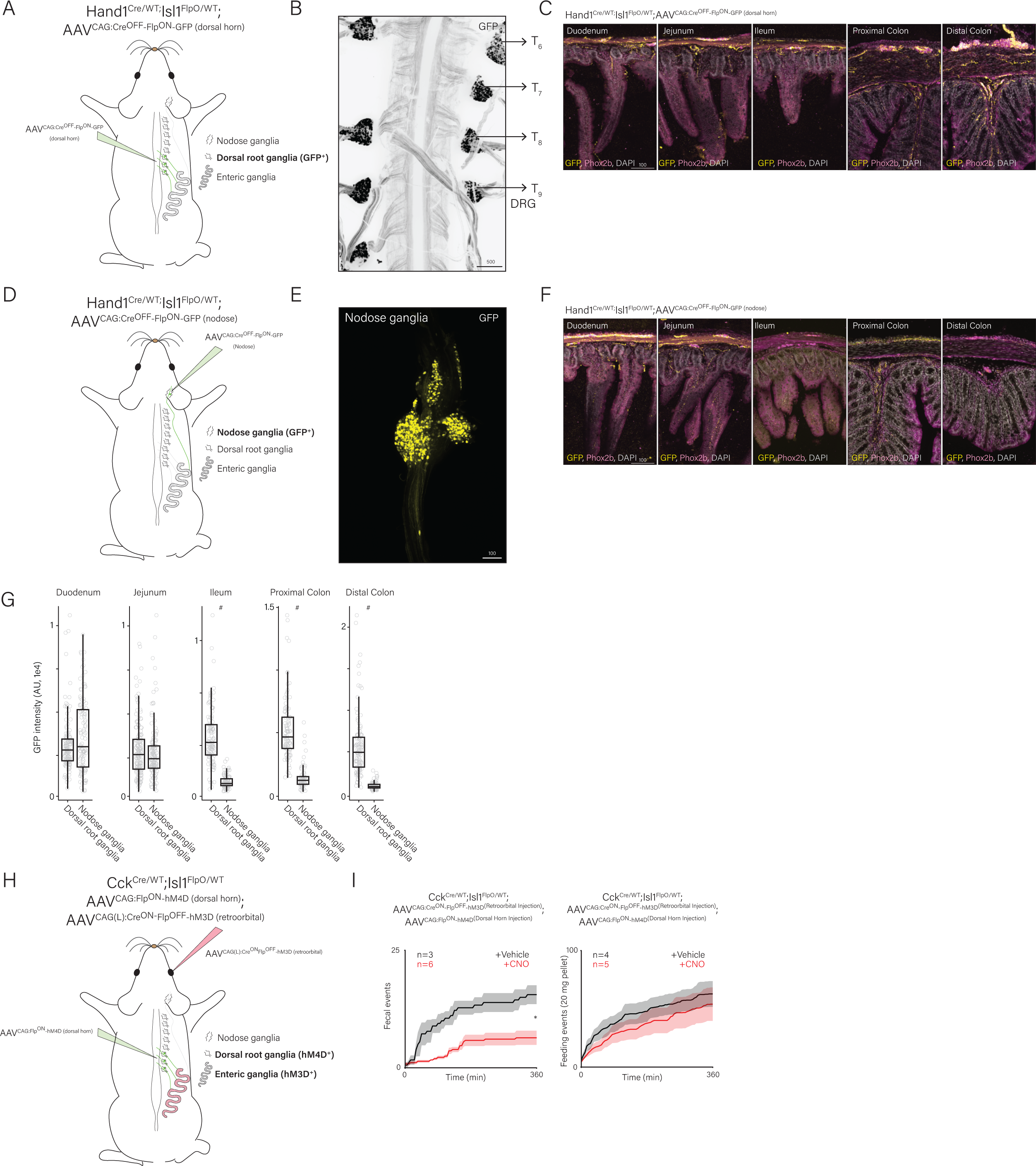
Extrinsic neurons broadly innervate the intestinal tract and relay signal from enteric neurons. **A)** Schematic representation of dorsal horn injection strategy used to isolate the sensory neurons from the dorsal root ganglia. **B)** Representative image of thoracic dorsal root ganglia transduced with GFP resulting from injection of CAG:Cre^OFF^-Flp^ON^-GFP into the dorsal horn of *Hand1^Cre^*;*Isl1^T2a-FlpO^* animals. Scale bar = 500µm. **C)** Representative confocal immunostaining images from *Hand1^Cre^*;*Isl1^T2a-FlpO^;CAG:Cre^OFF^-Flp^ON^-GFP^(Dorsal^ ^horn^ ^injections)^* animals in the indicated intestinal segment. *GFP* labeled axons from DRG sensory neurons are in yellow, *Phox2b^+^* enteric neurons are stained in magenta. Scale bar = 100µm. **D)** Schematic representation of direct nodose ganglia injection strategy used to isolate the sensory neurons from the nodose ganglia. **E)** Representative image of a whole mounted nodose ganglia transduced with GFP resulting from injection of CAG:Cre^OFF^-Flp^ON^-GFP into the nodose jugular complex of *Hand1^Cre^*;*Isl1^T2a-FlpO^*animals. Scale bar = 100µm. **F)** Representative confocal immunostaining images from *Hand1^Cre^*;*Isl1^T2a-FlpO^;CAG:Cre^OFF^-Flp^ON^-GFP^(nodose^ ^injections)^*animals in the indicated intestinal segment. *GFP* labeled axons from nodose ganglia sensory neurons are in yellow, Phox2b^+^ enteric neurons are stained in magenta. Scale bar = 100µm. **G)** Quantification of fluorescence intensity of axonal GFP in indicated segment of the intestine in *Hand1^Cre^;Isl1^FlpO^*animals with sensory neurons from (*Left*) dorsal root ganglia or (*Right*) nodose ganglia labeled. **H)** Schematic representation of viral transduction strategy to allow for simultaneous chemogenetic activation of the γ enteric neuron subclass ^(Cck-IRES-Cre;Isl1-T2a-FlpO;CAG:CreON-FlpOFF-hM3d)^ with concurrent silencing of thoracic dorsal root ganglia^(Cck-IRES-Cre;Isl1-T2a-FlpO;CAG:FlpON-hM4d;Dorsal horn injection)^ sensory neurons. **I)** *(Left)* Cumulative fecal production observed in response to vehicle or CNO injected treatment over six hours in *Cck^IRES-Cre^;Isl1^T2a-FlpO^;CAG:Cre^ON^-Flp^OFF^-hM3d;CAG:Flp^ON^-hM4d^(Dorsal^ ^horn^ ^injection)^* animals. Black traces represent the means for vehicle treated animals whereas red traces represent the means for CNO treated animals. Shaded areas represent mean ± SEM. *(Right)* Cumulative 20mg food pellet acquisition events recorded in overnight fasted animals treated with vehicle or CNO over six hours in *Cck^IRES-Cre^;Isl1^T2a-FlpO^;CAG:Cre^ON^-Flp^OFF^-hM3d;CAG:Flp^ON^-hM4d^(Dorsal^ ^horn^ ^injection)^* animals. Black traces represent the means for vehicle treated animals whereas red traces represent the means for CNO treated animals. Shaded areas represent mean ± SEM. *p<0.05 Wilcoxon-rank sum test done on total cumulative fecal pellets and cumulative food pellets retrieved, comparing vehicle or CNO injected animals. ^#^p<10^-3^ Wilcoxon-rank sum test.

We next sought to assess whether the projections of the dorsal root ganglia were indeed required for γ enteric neuron-mediated suppression of food acquisition. To test this, we transduced *Cck^IRES-Cre^;Isl1^T2a-FlpO^* animals with a CAG:Cre^ON^-Flp^OFF^-hM3D to enable precise CNO-based activation of γ enteric neurons. In these same animals, we also performed a dorsal horn injection of CAG:Flp^ON^-hM4D, so that we could enable CNO-based silencing^61^ of the dorsal root ganglia. We limited our AAV injections to transduce only the thoracic ganglia by restricting our injections to the thoracic dorsal horn segments. This location was chosen to minimize potential off-target effects of silencing non-intestine targeting DRG, as retrograde tracing studies indicate thoracic dorsal root ganglia represent the axial levels giving rise to ileal innervation^76^. In these animals, introduction of CNO would allow for simultaneous activation of γ enteric neurons and silencing of ileal-projecting dorsal root ganglia neurons. Consistent with a role for enteric-dorsal root ganglia communication, we observed that post-fasted food acquisition was only slightly reduced in CNO-treated *Cck^IRES-Cre^;Isl1^T2a-FlpO^; CAG:Cre^ON^-Flp^OFF^-hM3D;CAG:Flp^ON^-hM4D^(thoracic^ ^dorsal^ ^horn^ ^injection)^* animals compared to vehicle-treated controls. We also found that the reduction in these animals was significantly smaller compared to animals where γ enteric neurons were activated and the dorsal root ganglia pathways were unperturbed (**Figure 6G,H** compared to **Figure 4C**). Interestingly, we found that fecal output in CNO-treated *Cck^IRES-Cre^;Isl1^T2a-FlpO^; CAG:Cre^ON^-Flp^OFF^-hM3D;CAG:Flp^ON^-hM4D^(thoracic^ ^dorsal^ ^horn^ ^injection)^*animals was significantly reduced compared to vehicle-treated controls (**Figure 6H**) at a magnitude reminiscent of what we observed when only γ enteric neurons were activated (**Figure 4A**). This indicates that the dorsal root ganglia sensory pathway is not required for γ enteric neuron-mediated suppression of intestinal transit. Taken together, we propose that γ enteric neurons can relay signals to higher order neurons via dorsal root ganglia sensory neurons to modulate food consumption, whereas dorsal root ganglia signaling is not required for γ enteric neuron modulation of intestinal transit. Moreover, these data support a model in which the dorsal root ganglia and spinal cord, in addition to the nodose ganglia vagus nerve pathway, constitute a key gut-to-brain pathway critical for food intake.

## Discussion

Understanding neural control of intestinal functions requires phenotypic measurements in response to precise perturbation of clearly defined components in the intestinal circuitry. Here, we developed a flexible intersectional genetic system that allows for visualization and manipulation of the enteric neurons independently from extrinsic neurons. With this approach, we relate anatomical profiles of enteric neuron subclasses to defined genetic alleles, while further revealing remarkable subclass-specific effects on intestinal transit, fecal hydration, and food consumption. Interestingly, we find that enteric neuron control of food consumption is mediated, at least in part, by enteric-to-dorsal root ganglia signaling, expanding the roles of this understudied gut-to-brain pathway.

### Genetic access to enteric neuron subclasses

There is a long-standing precedent for defining enteric neuron subtype identity based on morphological, immunohistochemical, and electrophysiological features^1-5^. Many enteric neuron subtypes were initially characterized using the guinea pig and revealed striking heterogeneity in morphological organization, neurochemical content, and neurotransmitter identity. These studies, which have been extended to mice, rats, and humans, have resulted in a nuanced taxonomy of distinct enteric neuron cell types^1-6^. Broadly, the principal cell types include: the intrinsic primary afferent neurons, which relay signals from the villi to the broader enteric nervous system; interneurons, which connect enteric neurons to each other; excitatory and inhibitory motor neurons, which innervate smooth muscle; and secretomotor neurons, which control fluid balance. Identification of these cell types in tissue is challenging due to the required colocalization of multiple markers and anatomical features to triangulate neuronal identity. Amongst the growing catalog of markers, those more commonly used to define neural populations, such as *Chat*, *Nos1*, and *Calca (CGRP)*, are expressed in multiple intrinsic neuron populations and extrinsic neuron populations that project into the intestine^77-80^. As a result, mouse genetic alleles under the control of these and many other marker genes cause inadvertent alterations to multiple neuronal populations, limiting the ability to precisely assign functions to defined components in the broader intestinal circuit. We contend that it is important to disambiguate the contributions of enteric neurons subclasses from extrinsic neuronal influence. In this study, intersectional genetics facilitated the identification of enteric neuron subclasses that affect intestinal transit, fecal hydration, and even food consumption. Furthermore, the intersectional genetic tools allowed for the identification of an enteric-to-dorsal root ganglia pathway essential for food consumption, providing insight into a lesser studied gut-to-brain pathway. We propose that these considerations warrant, if not require, the continued use and development of intersectional genetic strategies to ensure sufficiently precise manipulation of defined neuronal populations.

### The roles of extrinsic ganglia in intestinal function

What are the roles of extrinsic projections in the intestine? As with enteric neurons, genetic isolation of extrinsic neuronal populations is challenging due to overlap of markers, but this can similarly be circumvented using the intersectional genetic strategies outlined here. Early efforts to characterize extrinsic innervation relied on combinations of histological analyses, pharmacological manipulations, and nerve stimulation studies. Surgical manipulations, such as chronic denervation or degradation of neural pathways into the intestine^64-66,81^, have had modest effects on intestinal peristalsis, which has been used to support the idea that the enteric nervous system is akin to a ‘second brain’^5^ with the ability to auto-regulate and function independently. However, this does not imply that extrinsic innervation is dispensable for a normally functioning bowel.

Of note is the extraordinarily expansive innervation by extrinsic neurons of the dorsal root^(sensory)^, nodose^(sensory)^, and sympathetic^(autonomic)^ ganglia of all parts of the intestine, ranging from the duodenum to the distal colon^29,30,76,82-88^, which is consistent with measurements made using the presented intersectional genetic strategies. Genetic separation of the two distinct sensory pathways (DRG vs Nodose) revealed that the dorsal root ganglia/spinal cord pathway most densely innervated the ileum and large intestine, which is consistent with recent anatomical observations^89^. By comparison, the nodose ganglia/vagus nerve most prominently innervated the upper bowel regions, namely the stomach and duodenum. Functionally, neurons in the nodose ganglia have been shown to discharge to low threshold or physiological intensities of stimulation^90-97^, whereas dorsal root ganglia neurons discharge to supraphysiological stimulus intensities^98-102^. These findings lend credence to the idea that dorsal root sensory afferents mediate nociception, while nodose afferents relay information related to internal state, such as satiety. However, recent studies with enhanced resolution have challenged this model by observing that sensory neurons in the dorsal root ganglia indeed respond to non-noxious stimuli in vivo^17,36^. A surge in the identification of extrinsic neuron subtypes has begun to reveal nuanced functions in multiple organ systems, particularly of sensory neurons^26,27,36,38,49,103-113^, although the functions of many extrinsic neuron remain to be characterized. Our study highlights an emergent role for the dorsal root ganglia sensory neurons in gut-to-brain sensory pathways, with select enteric neuron subclasses requiring intact dorsal root ganglia signaling for the regulation of food consumption.

### Interactions between the enteric nervous system and extrinsic ganglia

How do enteric and extrinsic neurons influence each other? In this study, independent labeling, and manipulation of enteric and extrinsic neurons, made possible by the mouse genetic system presented, enables insight into this question. Morphological labeling of distinct extrinsic populations confirmed that peripheral sensory and autonomic ganglia broadly innervate the intestine. Interestingly, we observed that the densest target of intestine-projecting extrinsic neurons was in fact enteric ganglia, whereas enteric neurons provided equally dense neural innervation to a broader range of targets within the intestine, such as other enteric ganglia, circular smooth muscle, and the villi. Further examination revealed that modulation of food consumption was most obvious for enteric neuron subclasses enriched in the ileum and proximal colon, and indeed the enteric neuron-dependent effects required intact communication with dorsal root ganglia sensory neurons. These observations support a model in which enteric neurons can initiate ascending signals via extrinsic sensory pathways to influence animal behavior. As an interesting example, the enteric nervous system of invertebrates was suggested to be critical for increases in maternal food intake during reproduction^114^. In addition to enteric-dorsal root ganglia signaling, it has also been suggested that a subpopulation of enteric neurons that send projections into sympathetic ganglia can also suppress appetite^115^. Future studies will be critical to uncover the different roles for sensory and autonomic neuron signaling as well as identification of distinct central circuits associated with these pathways. We propose that intersectional genetic approaches, such as those presented in this study, will be required to further elucidate these neural circuits.

Recent reports have found that extrinsic sensory innervation clearly exhibits mechano- and chemosensory roles in the small and large intestine^27,36,49,88,103,104,116-118^. Our observations support an extension of this model in which select subclasses of enteric neurons can influence food consumption via the dorsal root ganglia. It is tempting to speculate that the enteric ganglia may act as a ‘switchboard’, where the close apposition of extrinsic and intrinsic nervous systems allows for an environment where diverse neural lineages, namely enteric, sensory, and autonomic, can exert influence on each other’s activity and function. Consistent with this, subpopulations of neurons in each of the enteric, sensory, and autonomic lineage express neuropeptides, neurotransmitters, and receptors that could allow for pervasive crosstalk between all constituents of the intestinal neurocircuitry.

## Conclusions

In sum, we propose that further study of the enteric nervous system and broader intestine neural circuitry will require careful in vivo examination. The foundation for subsequent analyses will be critically dependent on the ability to precisely perturb defined components of the intestinal circuit in a robust and reproducible fashion. In this study, we have generated a broad array of mouse genetic tools that can be used in an intersectional manner to specifically target enteric neurons while ensuring that extrinsic neuronal populations remain unperturbed. We also couple these mouse genetic approaches with an adaptable AAV-based system that allows for Cre- and Flp-dependent reporters to be rapidly constructed. Taken together, these molecular genetic approaches have allowed for precise targeting of anatomically distinct enteric neuron subclasses, revealing distinct physiological consequences on intestinal function.

The emergence of single cell and single nuclei-based transcriptomics have greatly expanded the ability of peripheral neurobiologists to taxonomize neuronal subtypes based on gene expression. In fact, multiple efforts, both large- and small-scale, have been performed on peripheral neurons from the enteric^8,9,11,12,14,15^, sensory^10,16-27^ and autonomic^37-40^ lineages. The boundaries between cell types are difficult, perhaps even subjective, to define in a principled manner when relying primarily on transcriptomic data. Based on our anatomical and functional observations, it would be tempting to speculate that the α and β subclasses represent the excitatory and inhibitory circular muscle targeting motor neurons respectively; the γ, δ, and ζ subclasses represent interneurons; the η subclass represents secretomotor neurons; and the θ subclass represents intrinsic primary afferent neurons. This nomenclature must be flexible to future analysis that may identify non-trivial subtypes within each of these identities. We propose that subclass-specific recombinase driver lines presented here could be used ‘intersectionally’ with future subtype specific marker lines to target, manipulate, and examine populations defined in the future more precisely. Additionally, there certainly exist key populations of enteric neurons that have escaped capture by our current molecular genetic approaches, such as enteric intestinofugal neurons^42^ which are enteric neurons that send projections to sympathetic ganglia to create enteric-autonomic reflex loops. We suggest that enhancing the resolution of genome-wide approaches or implementing orthogonal epigenomic and cis-regulatory element profiling could lead to identification of currently inaccessible neuronal populations.

## Acknowledgements

We thank Chyuan-Sheng (Victor) Lin (Herbert Irving Comprehensive Cancer Center, Columbia University Medical Center) for key assistance with mouse model generation. We thank Michael Kissner (Columbia Stem Cell Initiative Flow Cytometry Core Facility, Columbia University Medical Center) for critical guidance in performing the FACS isolation. We thank David Ginty, Oliver Hobert, Josef Turecek, and Lynn Yap for critical discussions and comments on the manuscript. Parts of the schematics in Figure S5 were created with BioRender.com. This work was supported by a NIH 1DP2NS127278 (N.S.), The Whitehall Foundation (N.S.) and the Klingenstein-Simons Foundation (N.S.).

## Author contributions

D.S., P.R., C.J.W., and N.S. conceived, performed, and analyzed all aspects of the study. C.S., M.M., and N.S. designed, cloned, and tested the Cre and Flp dependent AAV constructs with help from other authors. D.S., P.R., C.J.W., and N.S. wrote the paper with input from all authors.

## Resource availability

Requests for resources and reagents should be directed to and will be fulfilled by the lead contact, Nikhil Sharma.

## Materials availability

Requests for mouse lines or recombinant DNA generated in this study should be directed to and will be fulfilled by Nikhil Sharma.

## Data and code availability

All data reported in this study will be shared by the lead contact upon request. All original code is available in this paper’s supplemental information. Any additional information required to reanalyze the data reported in this paper is available from the lead contact upon request.

## Methods

All experiments performed in this study were approved by the institutional animal care and use committee (IACUC) of Columbia University. Male and female mice used in this study were on the C57BL/6J background unless otherwise stated.

### Animals

The mouse lines presented in this study were generated using standard homologous recombination techniques in hybrid mouse embryonic stem cells. These mouse lines were generated by inserting a T2a-recombinase cassette directly upstream of the stop codon of the endogenous gene. This strategy was chosen to minimize disruption to endogenous gene expression and mitigate possible phenotypic consequences. Chimeras were generated by blastocyst injection and germline transmission confirmed by PCR using genomic DNA extracted from a tail biopsy. Unless stated otherwise, the neo selection cassette was left intact without obvious consequences. All mice generated in this study appear to be overtly indistinguishable from wild-type littermates, even when bred as homozygous carriers, and were born in Mendelian ratios. The mice generated in this study include: *Cox8b^T2a-FlpO^*, *Cys-ltr2^T2a-FlpO^*, *Elavl4^T2a-CreERT2^*, *Elavl4^T2a-FlpO^*, *Elavl4^T2a-FlpOERT2^*, *Hand1^T2a-FlpO^*, *Hmx3^T2a-CreERT2^*, *Hoxa5^T2a-CreERT2^*, *Hoxa5^T2a-FlpOERT2^*, *Hoxa7^T2a-FlpO^*, *Isl1^T2a-FlpO^*, *Kcns3^T2a-Cre^*, *Nxph2^T2a-Cre^*, *Rosa26:Cre^ON^-Flp^ON^-H2b^mTagRFP^*, *Rosa26:Cre^ON^-H2b^mTagRFP^*; *Rosa26:Flp^ON^-H2b^mTagRFP^*.

The tail-PCR genotyping strategies for the mice generated in this study are:

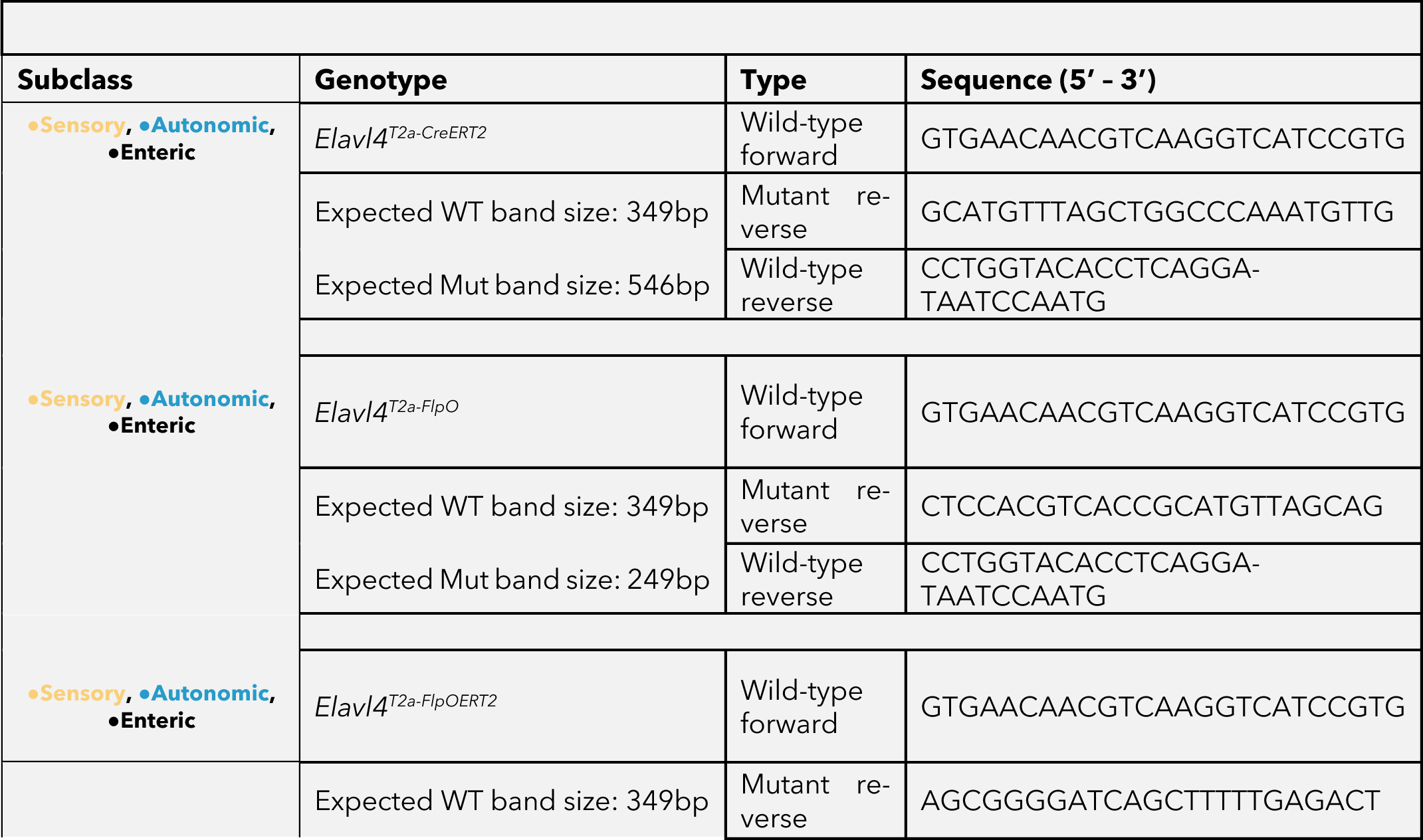
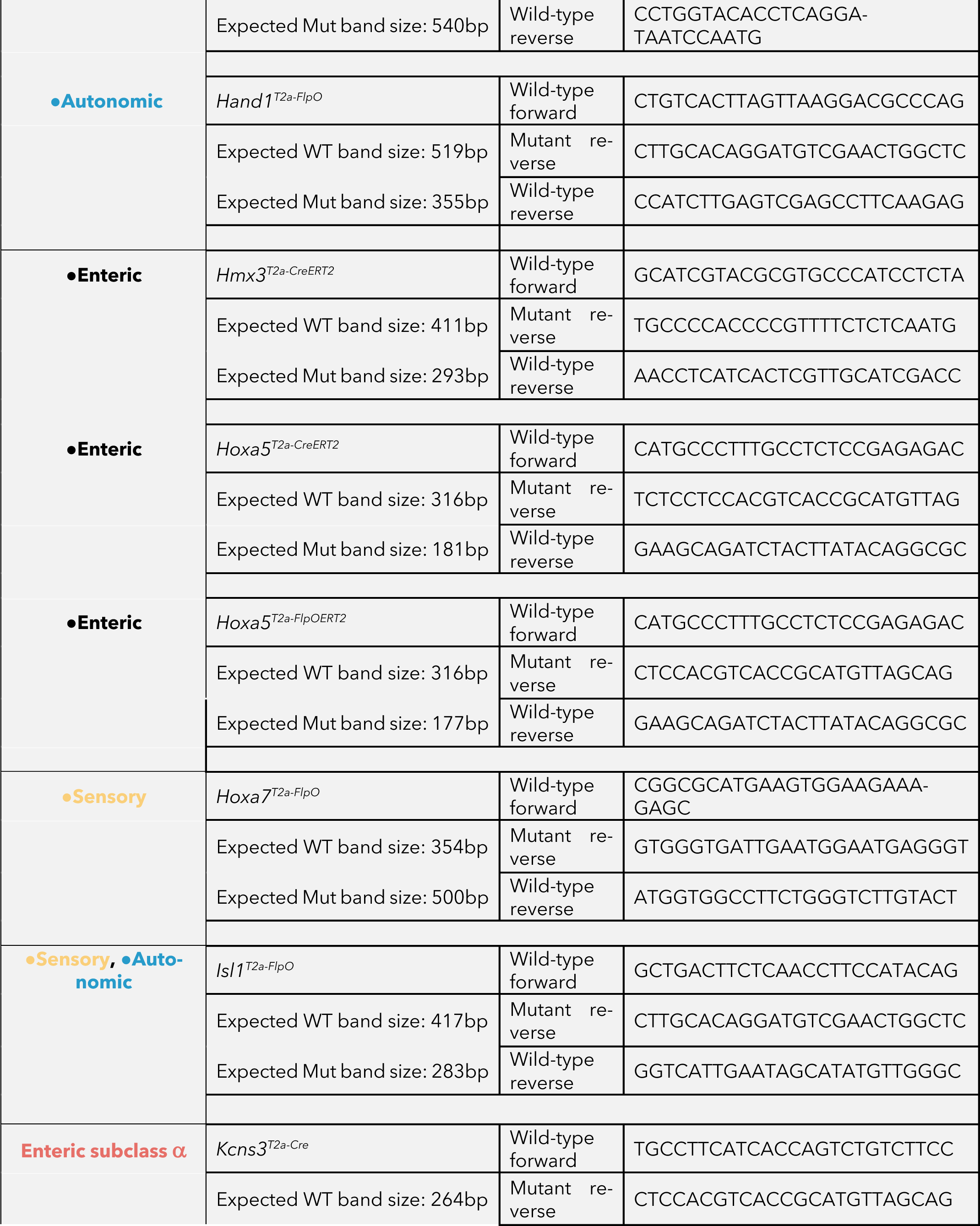
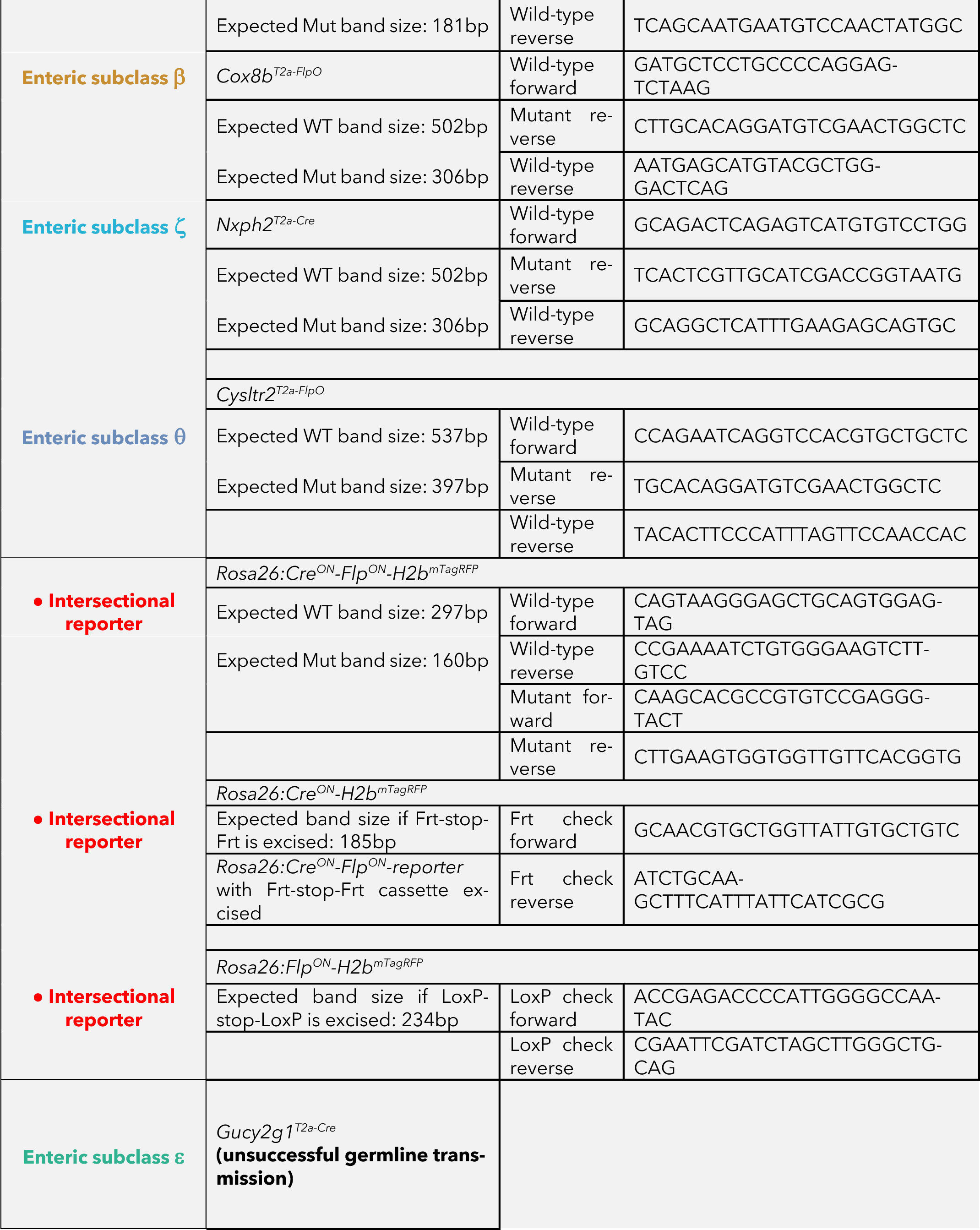

### Mouse generation

We designed all generated mouse lines by introducing *FlpO*, *FlpO^ERT2^*, and *Cre^ERT2^* knock-ins downstream of the marker gene of interest. We inserted a 2A autocatalytic peptide^55,56^ between the recombinase and the stop codon of the marker gene to minimize disruptions to endogenous gene expression (**Figure S1**). Rosa26 dual recombinase knock-in were generated using previously published strategies^119,120^. We note that introduction of Cre-dependent reporters into the *Kcns3^T2a-Cre^*allele did not lead to observable expression in the α-subclass of enteric neurons, and therefore was excluded from further analysis in this study. To generate lineage specific recombinase alleles, multiple independent strategies were attempted, with details for each described below:

For the peripheral enteric neuron lineage, three different strategies were attempted: Hmx3^T2a-CreERT2^;CAG:Cre^ON^-H2b^mTagGFP2^, Hoxa5^T2a-CreERT2^;CAG:Cre^ON^-H2b^mTagGFP2^, Hoxa5^T2a-FlpERT2^;CAG:Flp^ON^-H2b^mTagGFP2^, and Elavl4^T2a-CreERT2^;Isl1^T2a-FlpO^;CAG:Cre^ON^-Flp^OFF^-H2b^mTagGFP2^. While *H2b^mTagGFP2^* reporter expression was readily detected and restricted to enteric neurons in *Hmx3^T2a-CreERT2^*and *Elavl4^T2a-CreERT2^;Isl1^T2a-FlpO^* animals, we did not observe *H2b^mTagXFP^* reporter expression in any tissue examined *Hoxa7^T2a-CreERT2^* or *Hoxa7^T2a-FlpERT2^* animal. As a result, *Hoxa5^T2a-CreERT2^* and *Hoxa5^T2a-FlpERT2^* animals were excluded from further analysis in this study.

For the peripheral sensory neuron lineage, two different strategies were attempted: Hoxa7^T2a-FlpO^;CAG:Cre^ON^-H2b^mTagGFP2^ and Hand1^Cre^;Isl1^T2a-FlpO^;CAG:Cre^OFF^-Flp^ON^-H2b^mTagGFP2^. While *H2b^mTagGFP2^* reporter expression was readily detected and restricted to peripheral sensory ganglia in *Hand1^Cre^;Isl1^T2a-FlpO^*, we did not observe *H2b^mTagGFP2^* reporter expression in any tissue examined in *Hoxa7^T2a-FlpO^* animals. As a result, *Hoxa7^T2a-FlpO^*animals were excluded from further analysis in this study.

For the peripheral autonomic neuron lineage, one strategy was attempted: Hand1^T2a-FlpO^;CAG:Flp^ON^-H2b^mTagGFP2^. *H2b^mTagGFP2^* expression was readily detected and restricted to peripheral sympathetic ganglia in *Hand1^T2a-FlpO^*animals.

### Tamoxifen treatment

Tamoxifen was first dissolved in ethanol at a concentration of 20mg/ml then mixed with an equal volume of sunflower seed oil. 1:1 ethanol:sunflower seed oil was vigorously vortexed for 2-3 hours and then centrifuged under vacuum to fully remove residual ethanol. Single use aliquots of 20mg/ml tamoxifen in sunflower seed oil were stored at −80°C until needed. Indicated doses of tamoxifen were delivered by oral gavage in a total volume of ∼200µL between postnatal day 14-18. Generally, the dose of tamoxifen required for ‘dense’ labeling was 3-4mg, whereas for ‘sparse’ labeling a dose of 0.1-0.001mg was used.

### Single-cell RNA-sequencing preparation

Animals from the indicated genotypes were generally sacrificed after 7-14 days of reporter gene expression. Animals were first perfused with 1x PBS and then intestine was rapidly dissected and stored in ice-cold dissection media (DMEM:F12 media supplemented with penicillin-streptomycin). Segments from each region of the intestine, namely duodenum, jejunum, ileum, proximal and distal colon, were subdissected and rinsed to remove any food residue or fecal matter. Tissue segments were then cut up into shorter segments and pinned to a sylgard coated glass petri dish. An incision was made along the length of the intestine segment and insect pins were used to anchor the tissue such that the villi were facing up. Sharp forceps were used to separate the mucosal layer from the enteric plexuses. Isolated plexuses were stored in dissection media on ice for the remainder of the dissection period. Dissection periods were timed and were conducted for no longer than 2 hours to maximize tissue integrity. The remaining layers of the intestine were kept in a microfuge tube containing ice-cold DMEM:F12. Isolated plexuses were dissociated in 1mg/ml Liberase TH, 20 unit/mL Dispase, 0.1mg/ml DNAseI in DMEM:F12 + 1% pen/strep for 30 mins at 37°C. Digestion was quenched in media containing 20mg/mL BSA and 5mM EDTA in DMEM:F12, 1% pen/strep. Cell preparations were strictly maintained on ice/4°C for the remainder of the protocol. Digested tissue was gently triturated with fire-polished glass pipettes (opening diameter of approximately 100μm). Tissue was then passed through a 70-μm filter to remove cell doublets and debris. The neurons were pelleted and washed 4 times in 20 mg/ml BSA in DMEM:F12 + 1% penicillin-streptomycin followed by 2x washes with DMEM:F12 + 1% pen/strep all at 4°C. After washing, Tissue were resuspended in 45μl of DMEM:F12 + 1% Pen/strep, DyeCycle Ruby and Cytox Blue. Cells were sorted on a Sony MA-900 and gated on H2b^mTagXFP^ expression, DyeCycle Ruby and Cytox Blue signal to ensure intact and viable whole cells were collected. Sympathetic ganglia were processed in a similar fashion with the following modifications. Dissected sympathetic ganglia were first collected in DMEM:F12 supplemented with 1% pen/strep and 12.5 mM D-glucose. Dissociations were performed in 40 units papain, 4 mg/ml collagenase, 10 mg/ml BSA, 1 mg/ml hyaluronidase, 0.6 mg/ml DNase in DMEM:F12 + 1% pen/strep + 12.5 mM glucose for 30 min at 37°C. Digestion was quenched using 20 mg/ml ovomucoid (trypsin inhibitor), 20 mg/ml BSA in DMEM:F12 + 1% pen/strep + 12.5 mM glucose. Dissociated ganglia tissue were washed in a similar fashion to enteric plexus peels and resulting single-cell suspensions resuspended in an appropriate volume of DMEM:F12 + 1% pen/strep + 12.5 mM glucose. In all conditions, collected intact whole single cells were directly encapsulated for cDNA synthesis using the 10x Genomics platform.

### scRNA-seq analysis

Alignment, mapping, and general quality control were all conducted with the 10x Genomics Cell Ranger pipeline. This pipeline generated the gene expression tables for individual cells used in this study. All libraries were generated using the 10x Genomics Chromium Single Cell Kit v3. All samples were sequenced on a NextSeq 550 with 58bp sequenced into the 3’ end of the mRNAs. As quality control filter, individual cells were removed from the dataset if they had fewer than 1,000 discovered genes, fewer than 1,000 unique molecule identifiers (UMIs) or more than 10% of reads mapping to mitochondrial genes. Non-neuronal cells were identified in one of two ways. In the first, if the cells contained prominent markers smooth muscle or glia (*Acta2*, *Plp1*) they were excluded. In the second, if a cluster was clearly devoid of pan-neuronal marker genes (*Elavl4*, *Elavl3*), they were also excluded. Generally, we found cells isolated by FACS from Elavl4^T2a-FlpO^ animals were >95% neuronal in identity, cell isolated from Chat^IRES-Cre^ and Nos1^IRES-CreERT2^ mice were >85% neuronal in identity, and cells isolated from Baf53b^Cre^ animals were >70% neuronal in identity based on these criteria. Differential gene expression analysis was performed on all expressed genes using the FindMarker function in Seurat^121-125^ using the Wilcoxon-rank-sum test and a pseudocount of 0.001 was added to each gene to prevent infinite values. P values <10^−322^ were defined as 0, as the R environment does not handle numbers <10^−322^. Where appropriate, individual cell gene expression values were removed from the violin plots in the event inclusion would result in difficulty visualizing the data.

### AAV production and intravenous delivery

AAV vectors were designed with standard molecular biology and cloning techniques. Recombinant AAV was produced based on previously published protocols^126^. AAV genome plasmids were transfected in a mixture of pRC9, pHelper and the AAV-genome plasmid, or rAAV2-retro-helper, pHelper and the AAV-genome plasmid respectively into 12 T225 flasks of HEK 293T cells. Supernatant containing viral media were collected at 72 and 120 hours after transfection, and cells were scraped off and collected at 120 hours. AAV from cell pellets were extracted using a high salt lysis buffer containing 100unit/ml salt active nuclease in 40 mM Tris, 500 mM NaCl and 2 mM MgCl^2^ pH 8, referred to hereafter as SAN buffer. AAV from conditioned media was precipitated by via 8% PEG-800, 500mM NaCl induced precipitation. Precipitated virus was pelleted and resuspended in SAN buffer and combined with lysed cell pellets. Viral lysates were cleared by low-speed centrifugation (2000xg, 10 minutes, 4°C) then loaded onto an iodixanol density gradient for ultracentrifugation (60000rpm, 165 minutes, 18°C). Viral samples were then further concentrated using 100kD cutoff filter columns such that 12 T225 flasks resulted in approximately 50-60µL of concentrated virus. AAVs were stored in 1 × PBS supplemented with 0.01% Pluronic F-68, 35mM NaCl, 5% Glycerol. AAVs were stored in 10uL aliquots at −80°C until use. Viral gDNA was isolated for qPCR-based titer analysis by digestion in proteinase K. If viral titers were <10^14^ gc/ml, the viral preparation was discarded and not used in subsequent studies. 10^12^ genome copies of AAV in a total volume of 50-60µL were introduced intravenously by retroorbital injection to P13–15 mice. Before injection, all AAVs were supplemented with 1% fast green dye such that successful injection could be rapidly confirmed by the blueish appearance across the skin of animals.

### Immunohistochemistry

Animals were first perfused with 1x PBS and intestines rapidly removed and submerged in ice-cold dissection media (DMEM:F12 supplemented with 1% penicillin-streptomycin). Intestines were subdivided into duodenum, jejunum, ileum, proximal and distal colon and gently flushed with dissection media to remove undigested food particles and residual fecal matter. Cleaned intestines were dissected into 10-20mm segments and fixed in 1x PBS with 4% paraformaldehyde overnight at 4°C with end over end rotation. Tissues were then subsequently cryoprotected with 1x PBS with 30% sucrose for 12-18 hours at 4°C with end over end rotation. Ganglia tissue was processed in a similar fashion, with the dissection media being DMEM:F12 supplemented with 1% penicillin-streptomycin and 12.5mM D-Glucose. The tissue was embedded in NEG-50 mounting media and frozen at −80C until sectioning. Sections were acquired using a cryostat at a thickness of 40 microns and mounting on Superfrost plus slides. Tissue sections were stored at −80°C until used for immunostaining. For staining, slides were dried at room temperature for approximately one hour. Slides were rehydrated and washed with 1x PBS, 0.1% Triton X (IHC wash buffer) and then blocked in 5% normal donkey serum in IHC wash buffer for 60 minutes at 37°C. Primary antibody in wash buffer was added to slides and incubated overnight at 4°C followed by 3x 5-minute room temperature rinses with IHC wash buffer. Antibodies were visualized with Alexa-flour conjugated secondary antibodies applied to tissue sections in IHC wash buffer and incubated for 60 minutes at 37°C. Finally, samples were washed with IHC wash buffer supplemented with 0.001mg/mL DAPI to stain nuclei. Slides were mounted with a glycerol based mounting solution, sealed with nail polish and imaged on a W1-Yokogawa Spinning Disk Confocal within a week of staining.

### In situ hybridization

In situ hybridization (RNAscope) was performed largely following the recommended protocol from the manufacturer (https://acdbio.com/manual-assays-rnascope). In brief, relevant tissue was rapidly dissected from animals of the appropriate genotype, embedded in NEG-50 mounting media, and snap frozen in dry ice chilled 2-methylbutane. Cryosections at 10-15 micron thickness were obtained and slides stored at −80C until ready for use. Slides were fixed in prechilled 4% PFA in 1x PBS for 15 minutes at 4°C, then dehydrated using increasing concentrations of ethanol. Sections were pretreated with 5 minutes of protease III at room temperature. RNAscope probes were incubated with sections for 2 hours at 40°C followed by washes and addition of distinct fluorophores to visualize up to three different RNA targets. During the final washes, 0.001mg/ml DAPI was added to the wash buffer to visualize nuclei. Multichannel images were acquired using a W1-Yokogawa Spinning Disk Confocal and quantified using custom written macros in ImageJ and R.

### Behavioral measurements

All behavioral measurements were performed in custom built chambers intended to minimize experimental handling of the test animals. All behavioral measurements were fully automated and required minimal user interventions. Feeding studies were performed using a custom-built food pellet dispensing system with ad libitum access to drinking water. The specifications of the food pellet dispenser were constructed as described by the Feeding Experimentation Device (FED) 2.0 specifications^127,128^ (see https://hackaday.io/project/72964-feeding-experimentation-device-fed-20). Briefly, the housing of the dispenser was created using Stereolithography (SLA) 3D-printing in-house on a Formlabs Form 3+. Electronic components were acquired online, soldered and assembled together with the housing as per specifications. Finally, the code for pellet dispensing was flashed onto the Adafruit Feather M0 Adalogger microcontroller. Animals were fasted for approximately 12 hours prior to the start of the test period. Two 20mg food pellets and water were placed in the dispensing portion of the FED during the fasting period to ensure that mice could recognize food pellets dispensed from the FED. Food was then presented to the animal after the fast period and the number of 20 mg food pellet acquisitions were counted (see Figure S6 for schematic). All feeding assays commenced at 7:00 local time with food acquisition being continuously monitored for six hours. In rare cases, test animals were excluded if they acquired a large number of food pellets but did not eat them. For intestinal transit, animals were placed into a custom acrylic cut chamber with a wire-mesh flooring based on a previous published design^129^. Test animals were placed into these chambers at 7:00 local time with fecal samples falling through the wire-mesh flooring into a collection petri dish placed underneath the animals (see Figure S6 for schematic). To prevent fecal matter from adhering to the wire, food grade silicone lubricant was sprayed onto all surfaces prior to introducing animals into the test chambers. Fecal events were continually monitored for six hours and plotted as a function of time. Fluid content of fecal matter was determined largely as previously described^118^. Briefly, freshly produced fecal samples were immediately removed from the petri dish and weighed to determine the total weight. Fecal samples were then air dried for a week in chambers containing desiccant. After liquid content was removed, fecal samples were weighed again to determine dry weight. The dry weight divided by the total weight was defined as the hydration content of the fecal sample.

### Whole mount alkaline phosphatase staining of intestine

Sparse labeling and subsequent axonal arbor recovery of neurons was based on methods previously described for dorsal root ganglia sensory neurons^130^. For sparse labeling and axonal reconstructions of the α and β enteric neuron subclass, available CreER or FlpER lines were crossed to respective α and β selective recombinase driver lines. Density of labeling was controlled by the tamoxifen dose, with sparser labeling achieved by lower doses. 2-3 weeks after tamoxifen delivery, animals were perfused in PBS and intestinal tissue removed and placed in ice cold DMEM:F12 with 1% penicillin-streptomycin (dissection media). Intestinal tissue was flushed with dissection media to remove food particles and residual fecal mass. The tissue was then pinned to sylgard coated glass petri dishes. Spring scissors were used to open the intestine, with insect pins used to anchor the tissue onto the sylgard with the mucosal surface facing up. Fine forceps were used to remove the mucosal layer and expose the circular muscle and neuronal plexuses. Tissue was then fixed overnight at 4°C in 1x PBS with 4% PFA. The tissue was then washed in 1x PBS at room temperature and then incubated at 65-68°C for 2-2.5 hours to inactivate the endogenous alkaline phosphatase activity. The tissue was then washed with B3 buffer (0.1M Tris pH 9.5, 0.1M NaCl, 50mM MgCl2, 0.1% Tween-20) at room temperature. The substrate, NBT/BCIP (3.4µL per ml of B3 buffer), was added to start the reaction. After 2-3 hours at room temperature, tissue was washed in B3 buffer. In the event signal was not sufficiently developed, tissue was exposed to NBT/BCIP for an additional 2-3 hours at room temperature. In situations where neuronal morphology was readily detectable, images were acquired at this stage. Otherwise, the tissue was dehydrated by incubation in 50% ethanol (1 hour), then 75% ethanol (1hour), and lastly 100% ethanol overnight. After complete dehydration, tissue was cleared in BABB (Benzyl Alcohol: Benzyl Benzoate = 1:2) at room temperature for 30 mins. The tissue was then imaged under a Zeiss AxioZoom stereoscope. After imaging, the tissue was stored in 100% ethanol at 4°C. Single neuron reconstructions were generated in ImageJ by tracing axons emanating from the soma.

---

**Figure S1:**
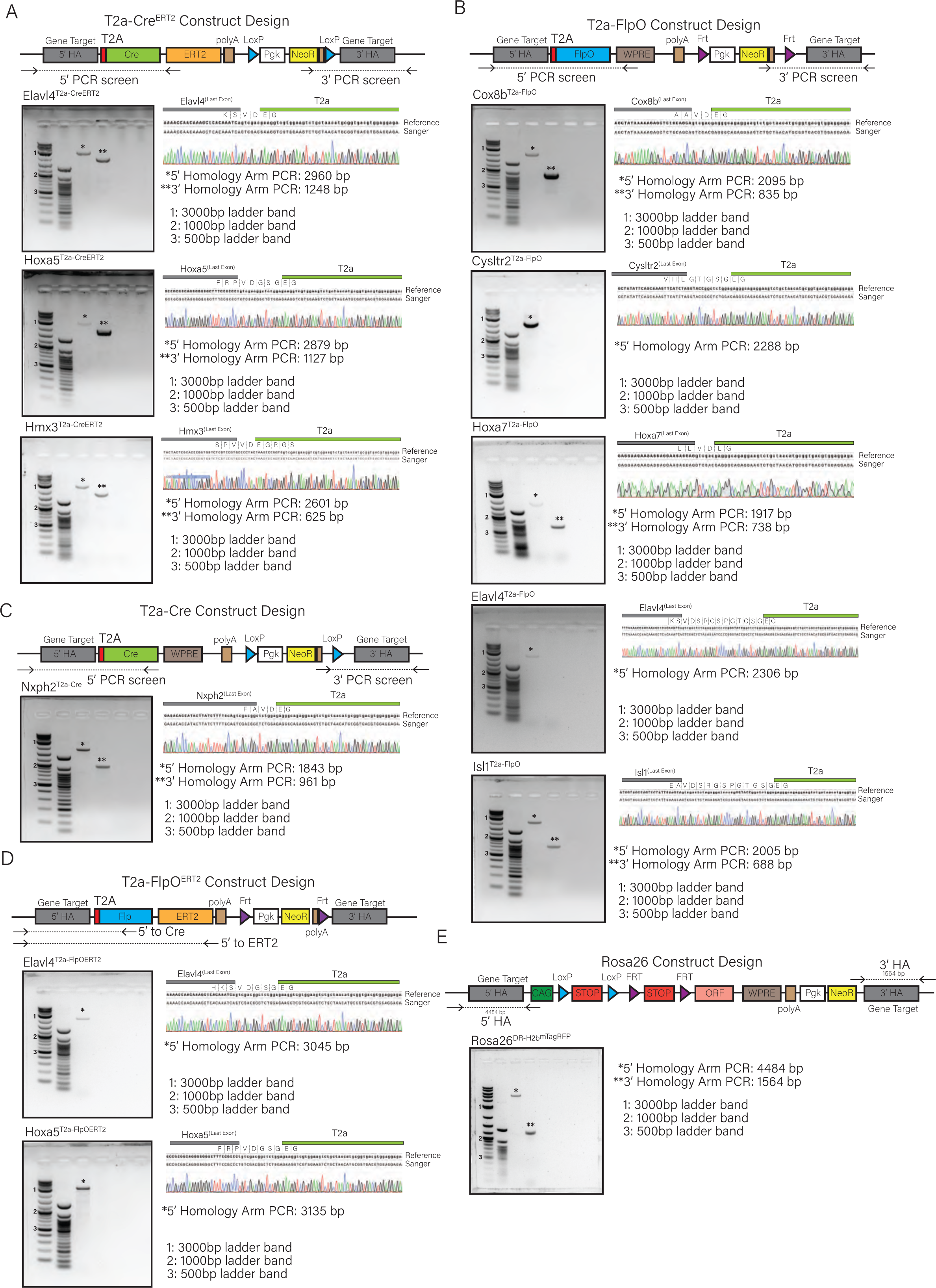
Gene targeting construct design and validation for recombinase and reporter driver alleles. **A)** *(Top)* Design strategy to introduce a T2a-Cre^ERT2^ cassette immediately upstream of the stop codon of the *Elavl4*, *Hoxa5*, and *Hmx3* loci. Schematic describing primer locations for the 5’ and 3’ long range PCR to confirm appropriate targeting to the endogenous locus. **B)** (Top) Design strategy to introduce a T2a-FlpO cassette immediately upstream of the stop codon of the *Cox8b*, *Cysltr2*, *Hoxa7*, *Elavl4* and *Isl1* loci. Schematic describing primer locations for the 5’ and 3’ long range PCR to confirm appropriate targeting to the endogenous locus. **C)** Design strategy to introduce a T2a-Cre cassette immediately upstream of the stop codon of the *Nxph2* locus. Schematic describing primer locations for the 5’ and 3’ long range PCR to confirm appropriate targeting to the endogenous locus. **D)** Design strategy to introduce a T2a-FlpO^ERT2^ cassette immediately upstream of the stop codon of the *Elavl4* and *Hoxa5* loci. Schematic describing primer locations for the 5’ and 3’ long range PCR to confirm appropriate targeting to the endogenous locus. **E)** Design strategy to introduce a lox-stop-lox-frt-stop-frt cassettes into the *Rosa26* locus. Schematic describing primer locations for the 5’ and 3’ long range PCR to confirm appropriate targeting to the *Rosa26* locus. Note that crossing these dual-recombinase dependent alleles to germline Cre or Flp recombinase deleter mice was used to generate lox-stop-lox and frt-stop-frt single recombinase dependent reporter alleles. In each case: *(Left)* Representative DNA agarose gel analysis for long-range PCR conducted with isolated genomic DNA from the indicated genotypes. *(Right)* Sanger sequencing analysis confirming correct targeting of the T2a-recombinase immediately upstream of the stop codon. Note that in select alleles the 3’ homology arm detection was unsuccessful. In these cases, 5’ PCR confirmation was extended to ensure that the entire T2a-recombinase-polyA-WPRE expression cassette was fully intact.

**Figure S2:**
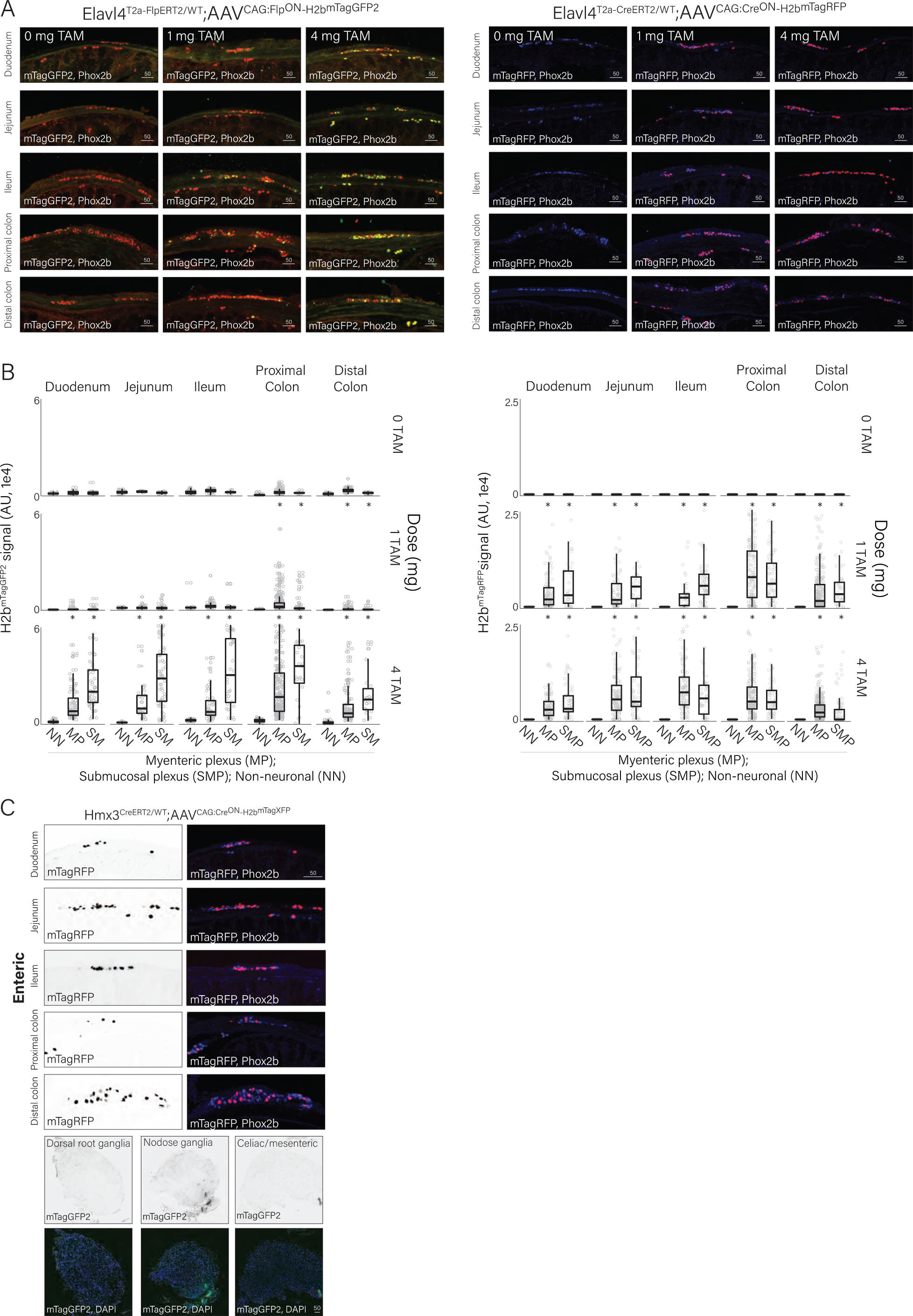
Validation of ligand inducible genetic strategies. **A)** Representative immunostaining images of tissue sections showing colocalization of reporter expression and the neuronal marker gene *Phox2b* in different regions of the intestine from (*Left*) *Elavl4^T2a-FlpOERT2^;CAG:Flp^ON^-H2b^mTagGFP2^*or (*Right*) *Elavl4^T2a-CreERT2^;CAG:Cre^ON^-H2b^mTagRFP^*animals in response to the indicated tamoxifen dose. Scale bar = 50µm. **B)** Quantification of *H2b^mTagXFP^*fluorescence intensity in neuronal and non-neuronal nuclei in (*Left*) *Elavl4^T2a-FlpOERT2^;CAG:Flp^ON^-H2b^mTagGFP2^* (*Right*) *Elavl4^T2a-CreERT2^;CAG:Cre^ON^-H2b^mTagRFP^* animals in the indicated tamoxifen dose. **C)** Representative immunostaining images of tissue sections showing colocalization of reporter expression and the neuronal marker gene Phox2b in (*Top*) different regions of the intestine or (*Bottom*) extrinsic ganglia from *Hmx3^T2a-CreERT2^;CAG:Cre^ON^-H2b^mTagRFP^*animals. Scale bar = 50µm. *p<10^-3^ Wilcoxon-rank sum test.

**Figure S3:**
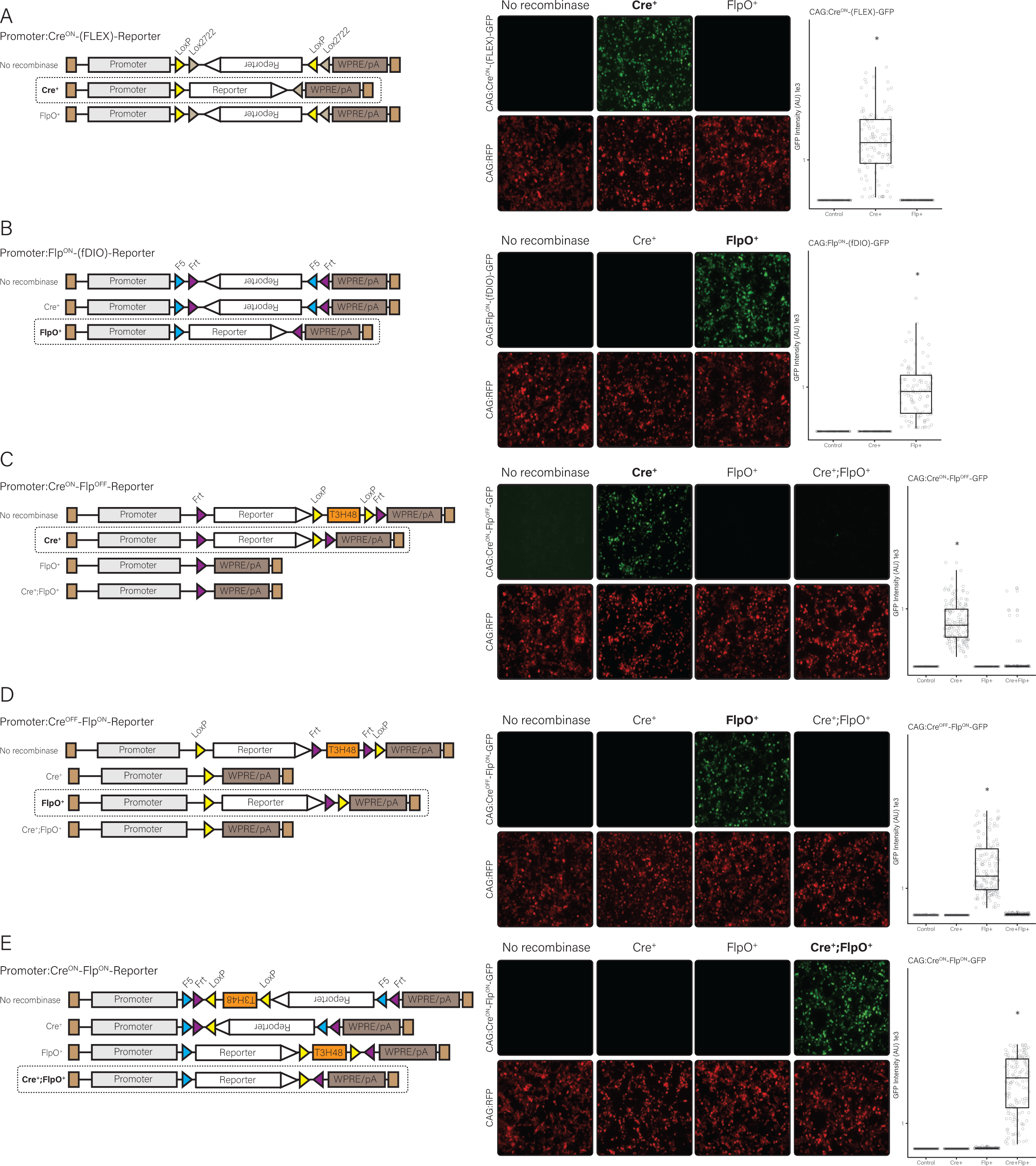
Generation of distinct recombinase dependent intersectional reporters constructs. **A)** *(Left)* Design of the Cre^ON^-reporter. *(Right)* Representative images show CAG:Cre^ON^-GFP expression in green and a constitutive CAG:RFP expression in red in 293T cells. **B)** *(Left)* Design of the Flp^ON^-reporter. *(Right)* Representative images show CAG:Flp^ON^-GFP expression in green and a constitutive CAG:RFP expression in red in 293T cells. **C)** *(Left)* Design of the Cre^ON^-Flp^OFF^-reporter. Expression is turned on by Cre only and turned off by Flp. *(Right)* Representative images show CAG:Cre^ON^-Flp^OFF^-GFP expression in green and a constitutive CAG:RFP expression in red in 293T cells. **D)** *(Left)* Design of the Cre^OFF^-Flp^ON^-reporter. Expression is turned on by Flp only and turned off by Cre. *(Right)* Representative images show CAG:Cre^OFF^-Flp^ON^-GFP expression in green and a constitutive CAG:RFP expression in red in 293T cells. **E)** *(Left)* Design of the Cre^ON^-Flp^ON^-reporter. Expression is turned on by both Cre and Flp only. *(Right)* Representative images show CAG:Cre^OFF^-Flp^ON^-GFP expression in green and a constitutive CAG:RFP expression in red in 293T cells. In all conditions, quantification of GFP intensities in indicated conditions are presented in the right most panels. *p<10^-3^ Wilcoxon-rank sum test.

**Figure S4:**
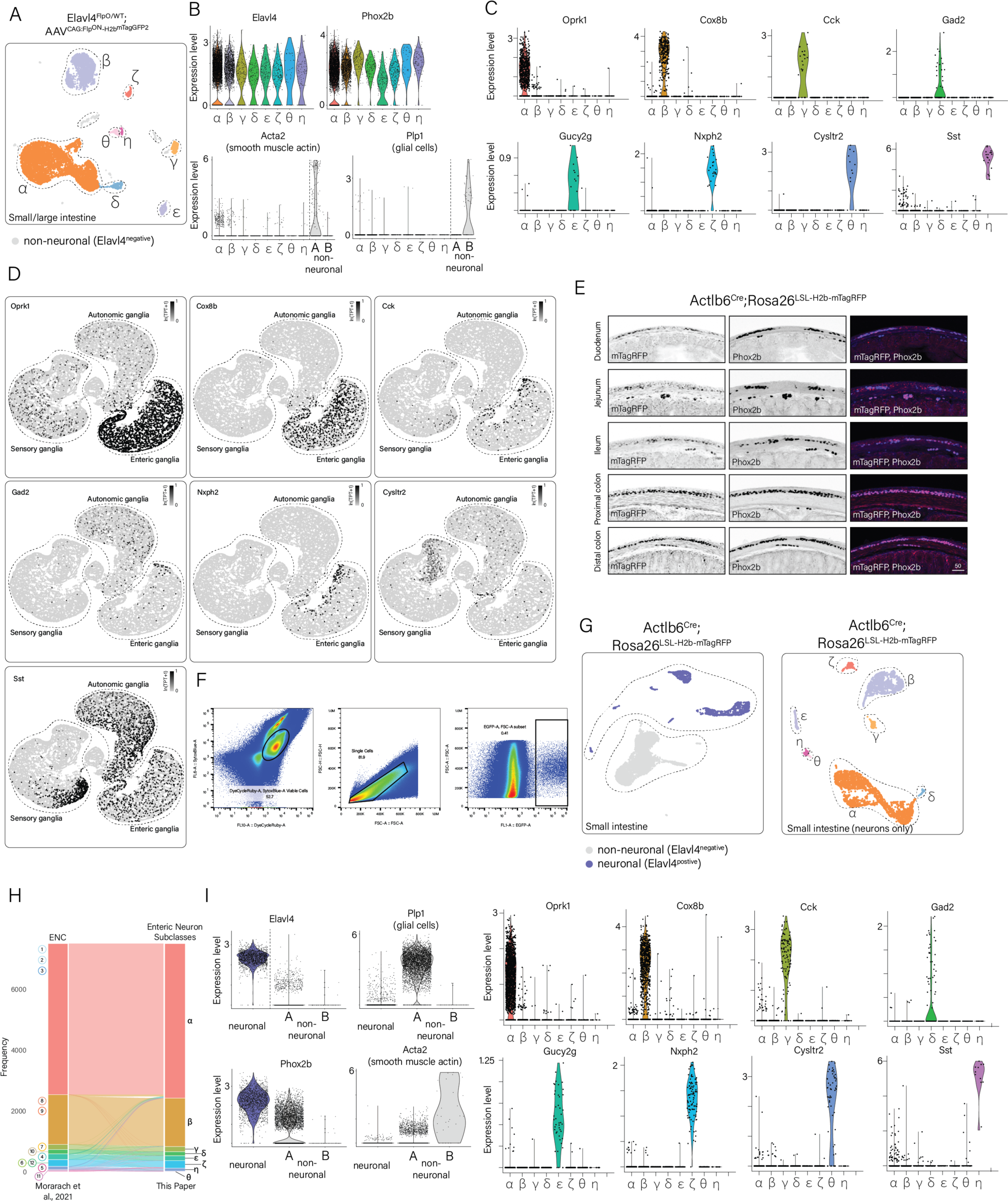
scRNA-seq analysis reveals enteric neuron subclass structure. **A)** UMAP visualizations of scRNA-seq data from *Elavl4^T2a-FlpO^;CAG:Flp^ON^-H2b^GFP^*mice clustered into transcriptomically distinct enteric neuron subclasses labeled by Greek letters, with α representing the cluster with the largest number of neurons, followed by β, etc. **B)** Violin plots displaying the expression of neuronal marker genes (*Elavl4, Phox2b*) and the lack of expression of non-neuronal marker genes (*Acta2, Plp1*) in enteric neuron subclasses derived from scRNA-seq data from *Elavl4^T2a-FlpO^;CAG: Flp^ON^-H2b^GFP^* mice. **C)** Violin plots displaying the transcriptomically distinct expression profiles for subclass-specific marker genes in enteric neuron subclasses derived from scRNA-seq data from *Elavl4^T2a-FlpO^;CAG: Flp^ON^-H2b^mTagGFP2^* mice. **D)** UMAP visualization showing enteric neuron subclass specific genes are often also expressed in peripheral sensory or autonomic neurons. **E)** Representative immunostaining images of tissue sections showing colocalization of reporter expression and the neuronal marker gene *Phox2b* in different regions of the intestine from *Actl6b^Cre^;Rosa26^LSL-H2b-mTagRFP^* mice. Scale bar = 50µm. **F)** Representative FACS plots with gating strategies for cell viability and fluorescence intensity. **G)** UMAP visualizations of enteric neuron scRNA-seq data from *Actl6b^Cre^;Rosa26^LSL-H2bRFP^*mice from the small intestine. (*Left)* UMAP visualization outlining the separation between neurons and non-neuronal cells. Neuronal clusters are identified by expression of *Elavl4* and *Phox2b*, non-neuronal clusters were identified by expression of *Acta2* and *Plp1*. (*Right*) Re-clustering of neurons with non-neuronal cells removed show a profile reminiscent to those derived from *Elavl4^T2a-FlpO^;CAG: Flp^ON^-H2b^GFP^* mice. **H)** Label transfer analysis between enteric neuron scRNA-seq data from Morarach et al and *Elavl4^T2a-FlpO^;CAG: Flp^ON^-H2b^GFP^* mice. **(I)** Violin plots displaying gene expression profiles of neurons and non-neuronal cells collected from Actl6b^Cre^; Rosa26^LSL-H2bRFP^ mice. *(Right)* Verification of *Elavl4*, *Hoxa5, Hmx3,* and *Phox2b* as enteric neuron markers. Non-neuronal cells primarily express markers like *Plp1* and *Acta2* associated with glia and smooth muscle cells. (*Left*) Violin plots displaying the transcriptomically distinct expression profiles for subclass-specific marker genes in enteric neuron subclasses derived from scRNA-seq data *Actl6b^Cre^; Rosa26^LSL-H2bRFP^*mice. TPT: tags per ten thousand.

**Figure S5:**
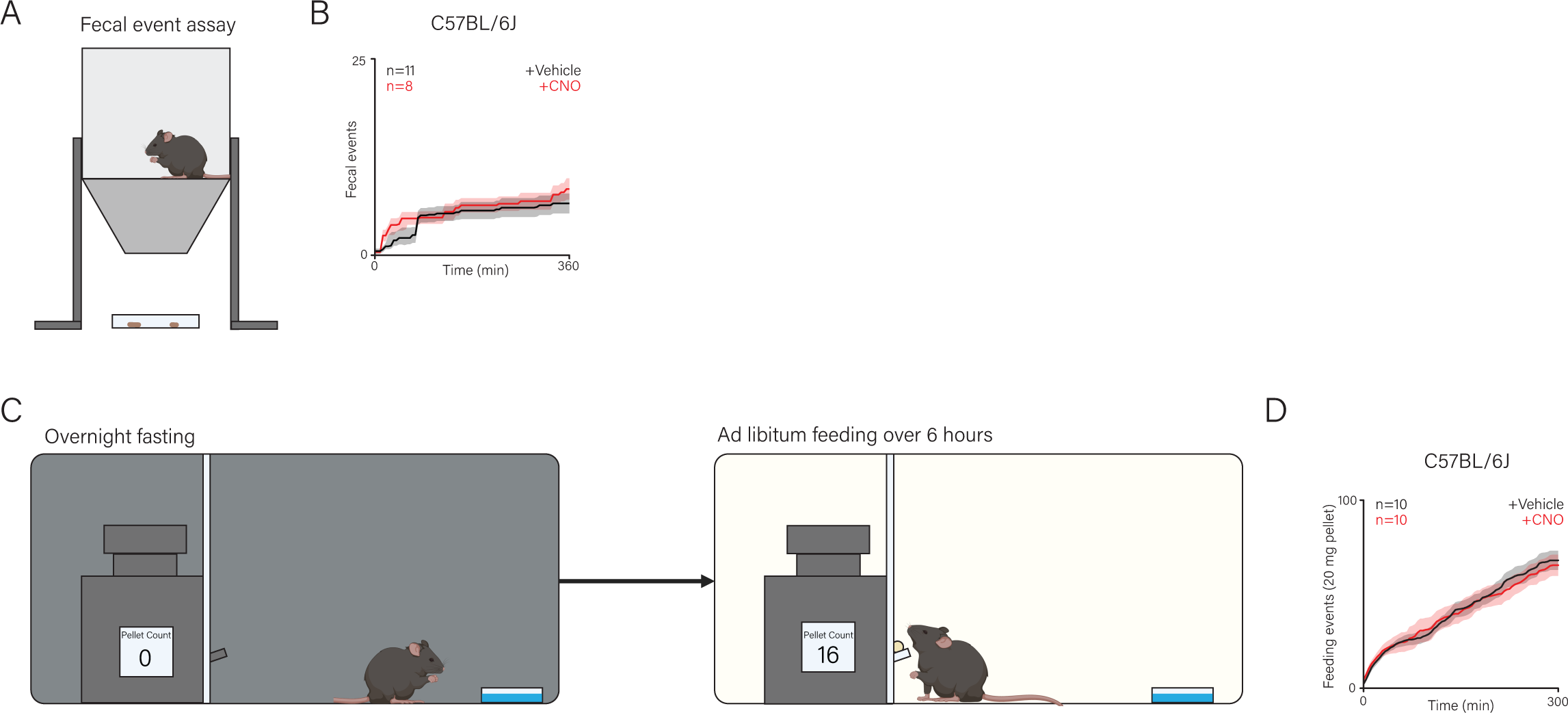
Behavior apparatus schematics for fecal production and food consumption. **A)** (*Left*) Schematic of the intestinal transit tracking assay. Animals are placed into the quiet chamber with a wire mesh flooring to allow for fecal samples to be collected or monitored rapidly without requiring manipulation by an experimenter. (*Right*) Cumulative fecal production over six hours observed in response to vehicle or CNO injected C57/Bl6 wild-type animals. Black traces represent the means for vehicle treated animals whereas red traces represent the means for CNO treated animals. Shaded areas represent mean ± standard error of the mean (SEM). **B)** (*Top*) Schematic of the food intake assay. Animals are fasted overnight in the test chamber. Subsequently, the automated feeder is activated, and the animal can acquire 20mg food pellets at will. Acquisition of food pellets is automated and does not require intervention by an experimenter. (*Bottom*) Cumulative 20mg food pellet acquisition events recorded in overnight fasted wild-type C57/Bl6 animals treated with vehicle or CNO over six hours. Black traces represent the means for vehicle treated animals whereas red traces represent the means for CNO treated animals. Shaded areas represent mean ± SEM. *p<0.05 Wilcoxon-rank sum test done on total cumulative fecal pellets and cumulative food pellets retrieved, comparing vehicle or CNO injected animals.

**Figure S6:**
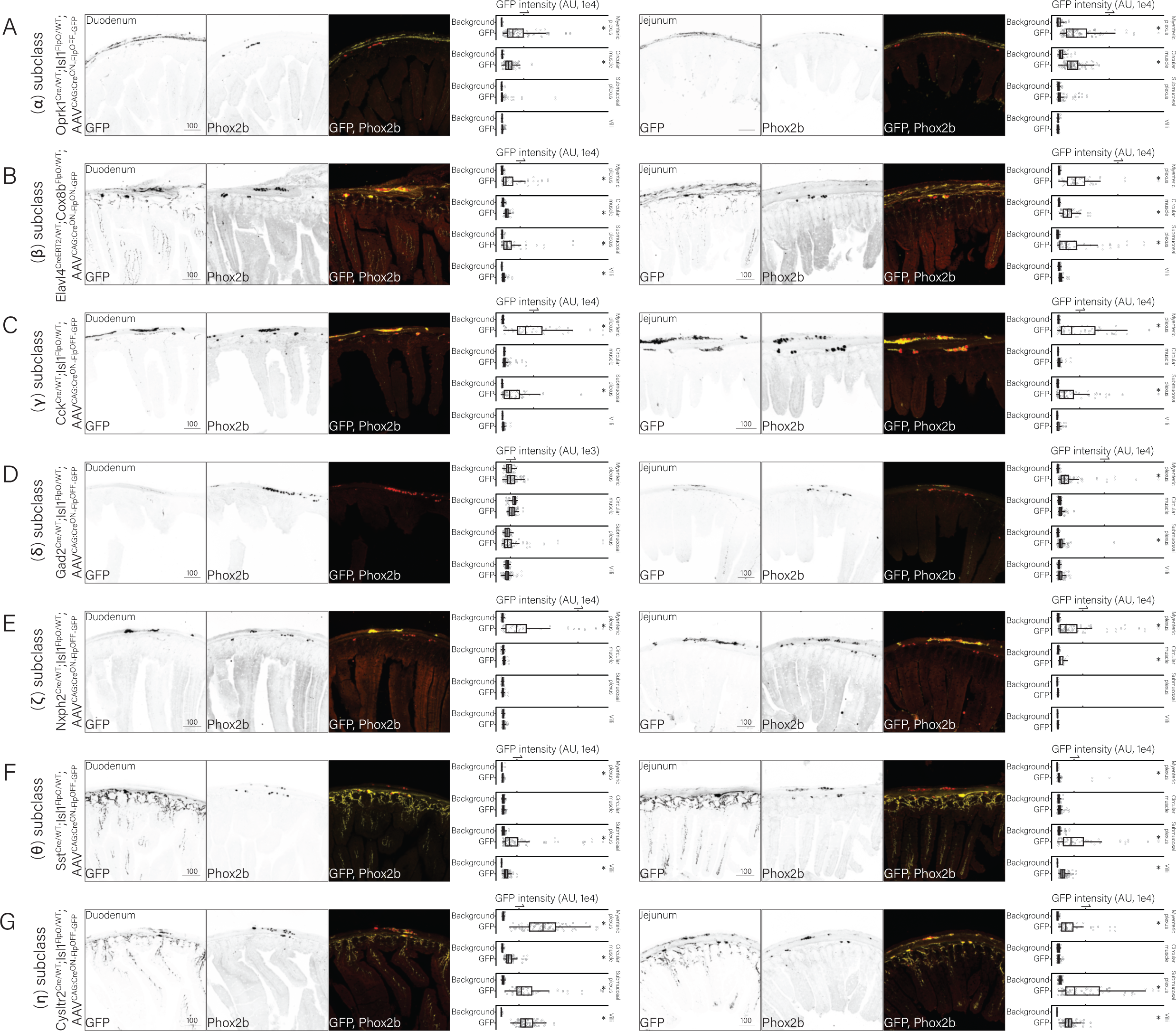
Genetically defined enteric neuron subclasses display distinct morphological profiles in the duodenum and jejunum. **A)** Analysis of fluorescence signal from enteric α subclass ^(*Oprk1-T2a-Cre;Isl1-T2a-FlpO*)^ using cytosolic GFP to reveal axonal arbors in the duodenum and jejunum. **B)** Analysis of fluorescence signal from enteric β subclass ^(*Elavl4-T2a-CreER;Cox8b-T2a-FlpO)*^ using cytosolic GFP to reveal axonal arbors in the duodenum and jejunum. **C)** Analysis of fluorescence signal from enteric γ subclass ^(*Cck-IRES-Cre;Isl1-T2a-FlpO*)^ using cytosolic GFP to reveal axonal arbors in the duodenum and jejunum. **D)** Analysis of fluorescence signal from enteric δ subclass ^(*Gad2-IRES-Cre;Isl1-T2a-FlpO*)^ using cytosolic GFP to reveal axonal arbors in the duodenum and jejunum. **E)** Analysis of fluorescence signal from enteric ζ subclass ^(*Nxph2-T2a-Cre;Isl1-T2a-FlpO*)^ using cytosolic GFP to reveal axonal arbors in the duodenum and jejunum. **F)** Analysis of fluorescence signal from enteric θ subclass ^(*Sst-IRES-Cre;Isl1-T2a-FlpO*)^ using cytosolic GFP to reveal axonal arbors in the duodenum and jejunum. **G)** Analysis of fluorescence signal from enteric η subclass ^(*Cysltr2-T2a-Cre;Isl1-T2a-FlpO*)^ using cytosolic GFP to reveal axonal arbors in the duodenum and jejunum. **(A-G)** *(Left)* Representative image of cytosolic *GFP*, the neuronal marker *Phox2b*, and an overlay in the duodenum or jejunum, *(Right)* Quantification of fluorescence intensity of cytosolic *GFP* in the duodenum or jejunum. *p<10^-3^ Wilcoxon-rank sum test. Scale bar for cytosolic GFP/Phox2b images = 100µm.

**Figure S7:**
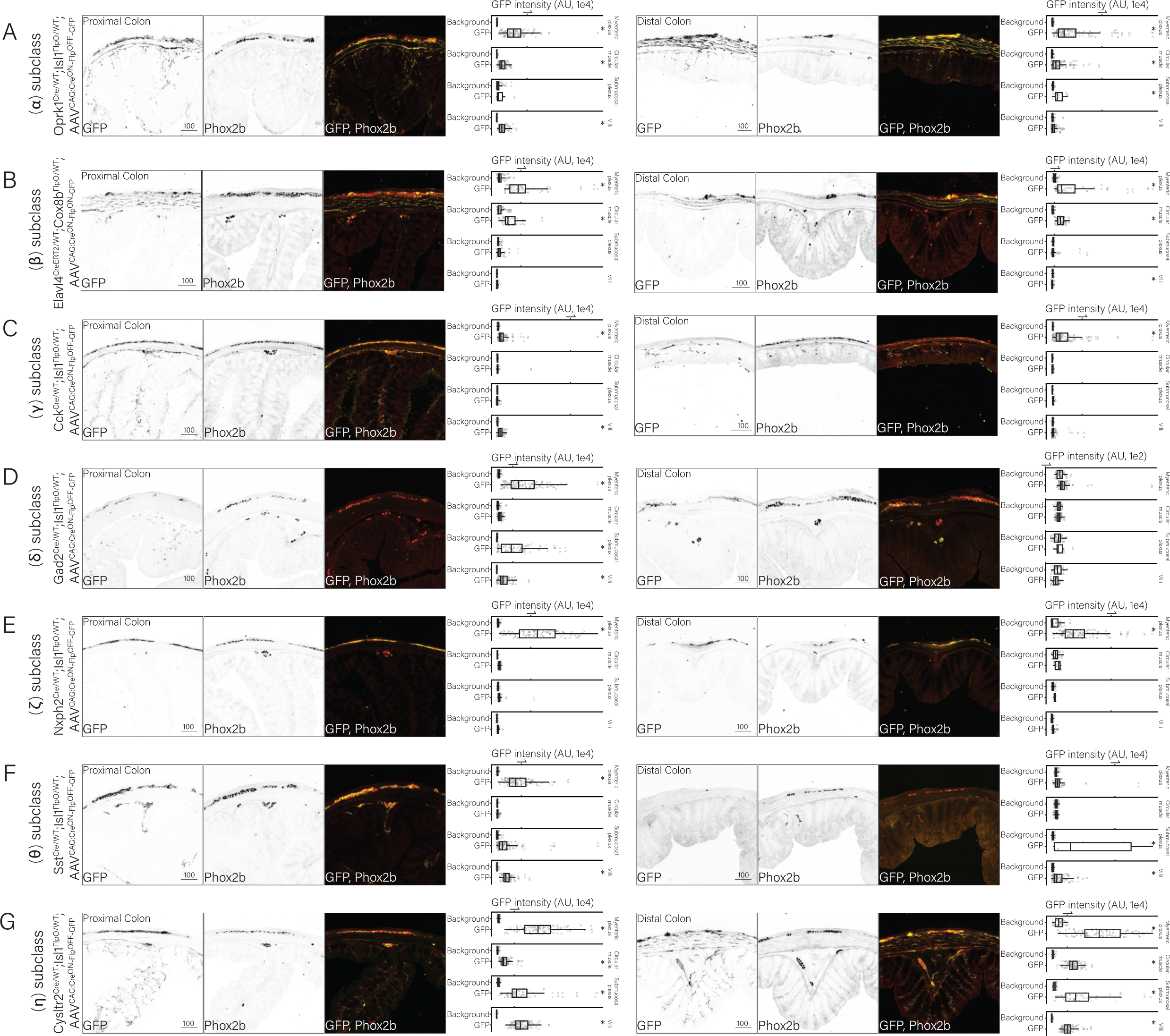
Genetically defined enteric neuron subclasses display distinct morphological profiles in the proximal and distal colon. **A)** Analysis of fluorescence signal from enteric α subclass ^(*Oprk1-T2a-Cre;Isl1-T2a-FlpO*)^ using cytosolic GFP to reveal axonal arbors in the proximal and distal colon. **B)** Analysis of fluorescence signal from enteric β subclass ^(*Elavl4-T2a-CreER;Cox8b-T2a-FlpO*^ using cytosolic GFP to reveal axonal arbors in the proximal and distal colon. **C)** Analysis of fluorescence signal from enteric γ subclass ^(*Cck-IRES-Cre;Isl1-T2a-FlpO*)^ using cytosolic GFP to reveal axonal arbors in the proximal and distal colon. **D)** Analysis of fluorescence signal from enteric δ subclass ^(*Gad2-IRES-Cre;Isl1-T2a-FlpO*)^ using cytosolic GFP to reveal axonal arbors in the proximal and distal colon. **E)** Analysis of fluorescence signal from enteric ζ subclass ^(*Nxph2-T2a-Cre;Isl1-T2a-FlpO*)^ using cytosolic GFP to reveal axonal arbors in the proximal and distal colon. **F)** Analysis of fluorescence signal from enteric θ subclass ^(*Sst-IRES-Cre;Isl1-T2a-FlpO*)^ using cytosolic GFP to reveal axonal arbors in the proximal and distal colon. **G)** Analysis of fluorescence signal from enteric η subclass ^(*Cysltr2-T2a-Cre;Isl1-T2a-FlpO*)^ using cytosolic GFP to reveal axonal arbors in the proximal and distal colon. **(A-G)** *(Left)* Representative image of cytosolic *GFP*, the neuronal marker *Phox2b*, and an overlay in the proximal or distal colon, *(Right)* Quantification of fluorescence intensity of cytosolic *GFP* in the proximal or distal colon. *p<10^-3^ Wilcoxon-rank sum test. Scale bar for cytosolic GFP/Phox2b images = 100 µm.

**Figure S8:**
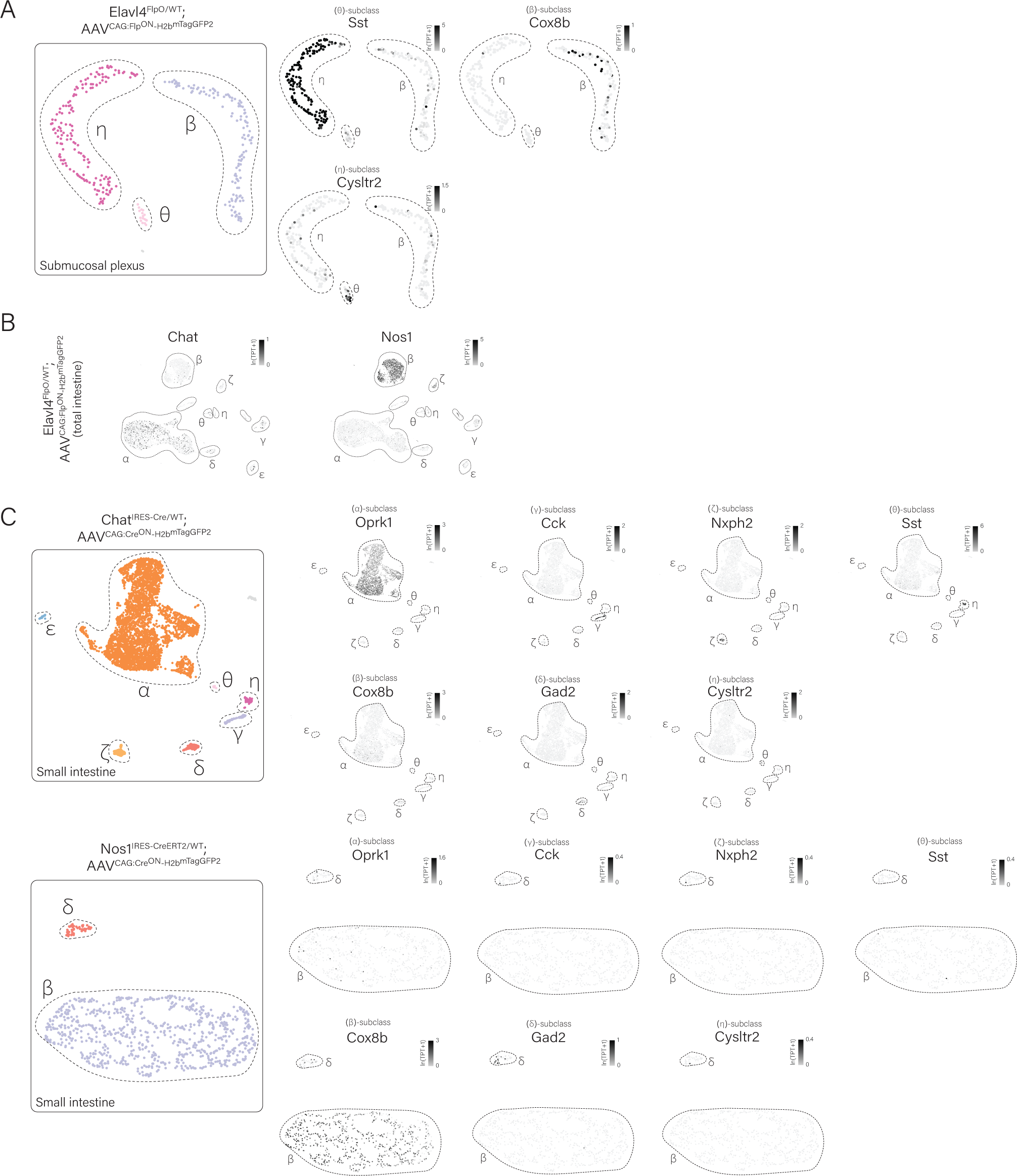
Iterative scRNA-seq reveals neurochemical features of the genetically defined enteric neuron subclasses. **A)** UMAP visualizations of enteric neuron scRNA-seq data derived from intact whole enteric neurons collected by FACS from the submucosal plexus using *Elavl4^T2a-FlpO^;CAG:Flp^ON^-H2b^mTagGFP2^* mice. **B)** UMAP visualizations of (*Left*) *Chat* and (*Right*) *Nos1* expression in enteric neuron subclass clusters FACS isolated from *Elavl4^T2a-FlpO^;CAG:Flp^ON^-H2b^mTagGFP2^*mice. **C)** (*Left*) UMAP visualizations of enteric neuron scRNA-seq data derived from intact whole enteric neurons collected by FACS from *Chat^IRES-Cre^;CAG:Cre^ON^-H2b^mTagGFP2^*mice. (*Right*) UMAP visualizations of subclass-specific gene markers expression in the selected dataset. **D)** UMAP visualizations of enteric neuron scRNA-seq data derived from intact whole enteric neurons collected by FACS from *Nos1^IRES-CreERT2^;CAG:Cre^ON^-H2b^mTagGFP2^*mice. (*Right*) UMAP visualizations of subclass-specific gene markers expression in the selected dataset. TPT: tags per ten thousand.

**Figure S9.**
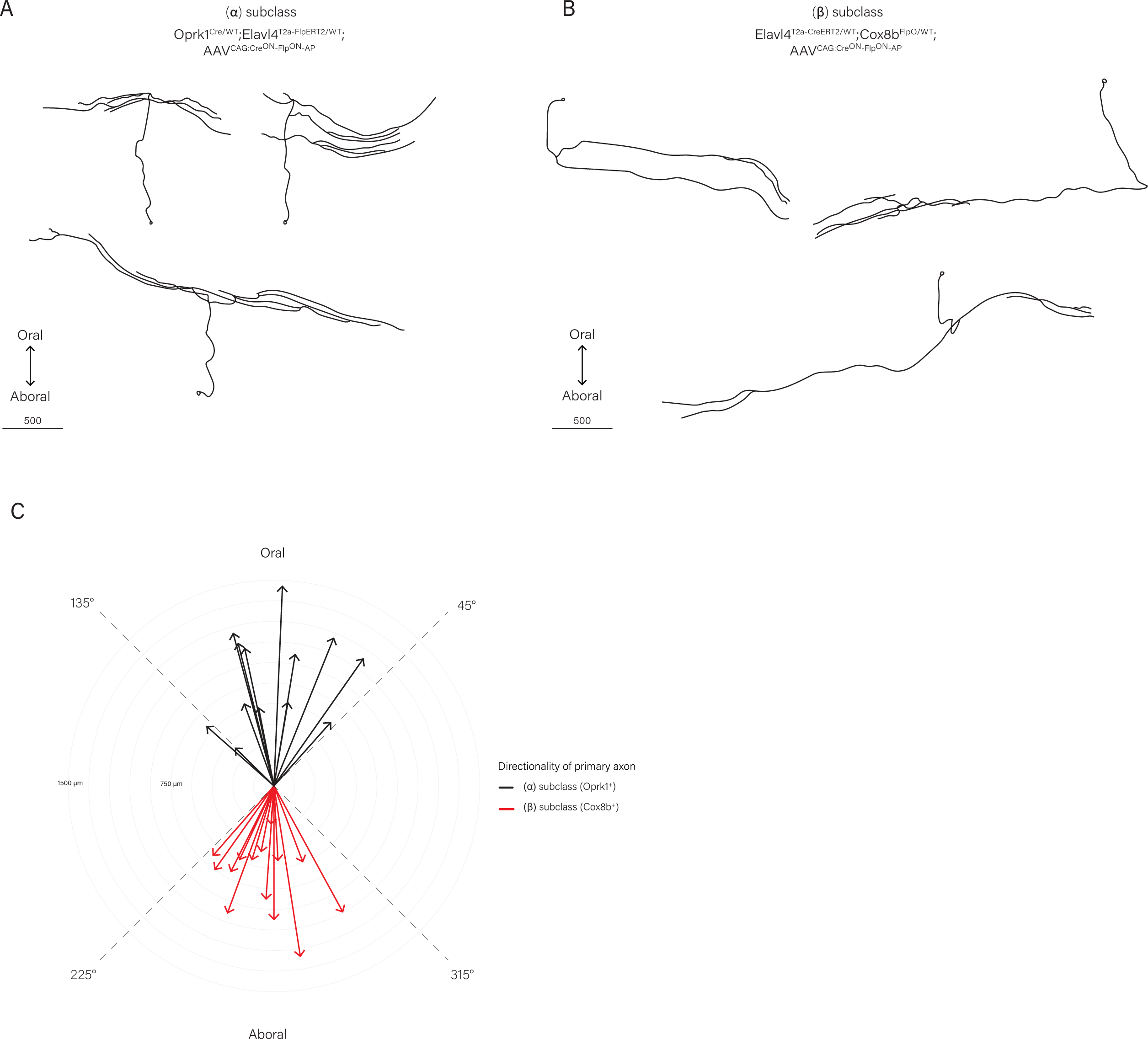
α & β enteric neuron populations display polarized projections. A) Representative reconstructions of axonal arbors using AP staining with tissue collected from *(Left)* α^(*Oprk1-T2a-Cre;Isl1-T2a-FlpO*)^;CAG:Cre^ON^-Flp^ON^-AP. Oral-aboral orientation is noted. Scale bar = 500 µm. B) Representative reconstructions of axonal arbors using AP staining with tissue collected from β^(*Elavl4-T2a-CreER;Cox8b-T2a-FlpO*)^;CAG:Cre^ON^-Flp^ON^-AP mice. Oral-aboral orientation is noted. Scale bar = 500 µm. **B)** Quantification of axonal projection directionality for individual reconstruction of neurons from the α & β enteric neuronal populations displayed on a polar coordinate system. Each arc represents an increment of 150 µm.

